# Relative pitch representations and invariance to timbre

**DOI:** 10.1101/2022.01.13.476197

**Authors:** Malinda J. McPherson, Josh H. McDermott

**Author notes:** Corresponding author, MIT, Building 46-4078c, 77 Massachusetts Avenue, Cambridge, MA, 02139.

## Abstract

Information in speech and music is often conveyed through changes in fundamental frequency (f0), perceived by humans as “relative pitch”. Relative pitch judgments are complicated by two facts. First, sounds can simultaneously vary in timbre due to filtering imposed by a vocal tract or instrument body. Second, relative pitch can be extracted in two ways: by measuring changes in constituent frequency components from one sound to another, or by estimating the f0 of each sound and comparing the estimates. We examined the effects of timbral differences on relative pitch judgments, and whether any invariance to timbre depends on whether judgments are based on constituent frequencies or their f0. Listeners performed up/down and interval discrimination tasks with pairs of spoken vowels, instrument notes, or synthetic tones, synthesized to be either harmonic or inharmonic. Inharmonic sounds lack a well-defined f0, such that relative pitch must be extracted from changes in individual frequencies. Pitch judgments were less accurate when vowels/instruments were different compared to when they were the same, and were biased by the associated timbre differences. However, this bias was similar for harmonic and inharmonic sounds, and was observed even in conditions where judgments of harmonic sounds were based on f0 representations. Relative pitch judgments are thus not invariant to timbre, even when timbral variation is naturalistic, and when such judgments are based on representations of f0.

## 1 INTRODUCTION

A central challenge for our perceptual systems is that we must often make judgments about one variable amid variation across other variables (Carruthers et al., 2015; DiCarlo & Cox, 2007; Liu et al., 2019; Sharpee et al., 2011). For example, object shape must be estimated across changes in lighting and pose, and spoken words recognized across variation in the voice of the speaker, manner of speaking, and listening environment.

Another instance of this challenge can be found in pitch perception. ‘Pitch’ can broadly refer to the attribute that allows us to order sounds from low to high (ANSI, 1994), but has traditionally been defined within hearing research as the perceptual correlate of a sound’s fundamental frequency (f0) (Plack et al., 2005). Natural sounds are often harmonic, such that their constituent frequencies (called harmonics, or partials) are integer multiples of a common f0. In some contexts, such as voice recognition, the absolute f0 of a sound is perceptually important (McPherson & McDermott, 2018). The ability to estimate and make judgments about the absolute f0 is known as absolute pitch (not to be confused with the ability to name musical notes, colloquially known as ‘perfect pitch’, which is sometimes also referred to as absolute pitch in the scientific literature). But many of the judgments we must make involve comparisons between sounds. For instance, we often need to determine if a prosodic contour in speech ascended or descended, or if the pitch intervals between the notes of a melody match those in memory. Such decisions about the direction and magnitude of pitch changes are known as relative pitch. One way to make relative pitch judgments is to estimate the f0 of individual sounds and then compare the estimates (McPherson et al., 2022; McPherson & McDermott, 2018; McPherson & McDermott, 2020; Micheyl et al., 2010; Moore & Glasberg, 1990). However, when the f0 changes between two sounds, there are corresponding changes in the frequencies of individual harmonics, which shift up or down with the f0. In many conditions listeners appear to use a representation of these shifts between constituent frequency partials (Demany & Ramos, 2005), rather than representations of f0, to judge whether one sound is higher or lower than another (Faulkner, 1985; McPherson et al., 2022; McPherson & McDermott, 2018; McPherson & McDermott, 2020). Relative pitch can thus be extracted in two different ways. Listeners use representations of f0 when sounds are presented in noise, or must be remembered over time, but use representations of frequency partials when sounds are presented back-to-back in quiet conditions.

Invariance presents a challenge for pitch perception because the relative amplitudes of harmonics can vary between sounds that must be compared, such that sounds with the same f0 can have different spectral envelopes (Figure 1a). This variation is intrinsic to the source-filter generative model by which speech and many instrument sounds are produced (Fletcher & Rossing, 2010; Stevens, 2000). The source is characterized in part by its f0. The filter is mediated by the vocal tract or instrument body, which have resonances that amplify some frequencies and attenuate others. In acoustic terms, the filter alters the spectral envelope of a sound. In perceptual terms, it alters a sound’s timbre. The computational challenge is that the auditory system receives the aggregate sound spectrum (as transduced by the cochlea) and from this must derive both pitch and timbre. Pitch perception is nonetheless believed to be somewhat invariant to the variation imposed by filtering in the sound generation process (de Cheveigne, 2010; Demany & Semal, 1993; Semal & Demany, 1991).

**Figure 1:**
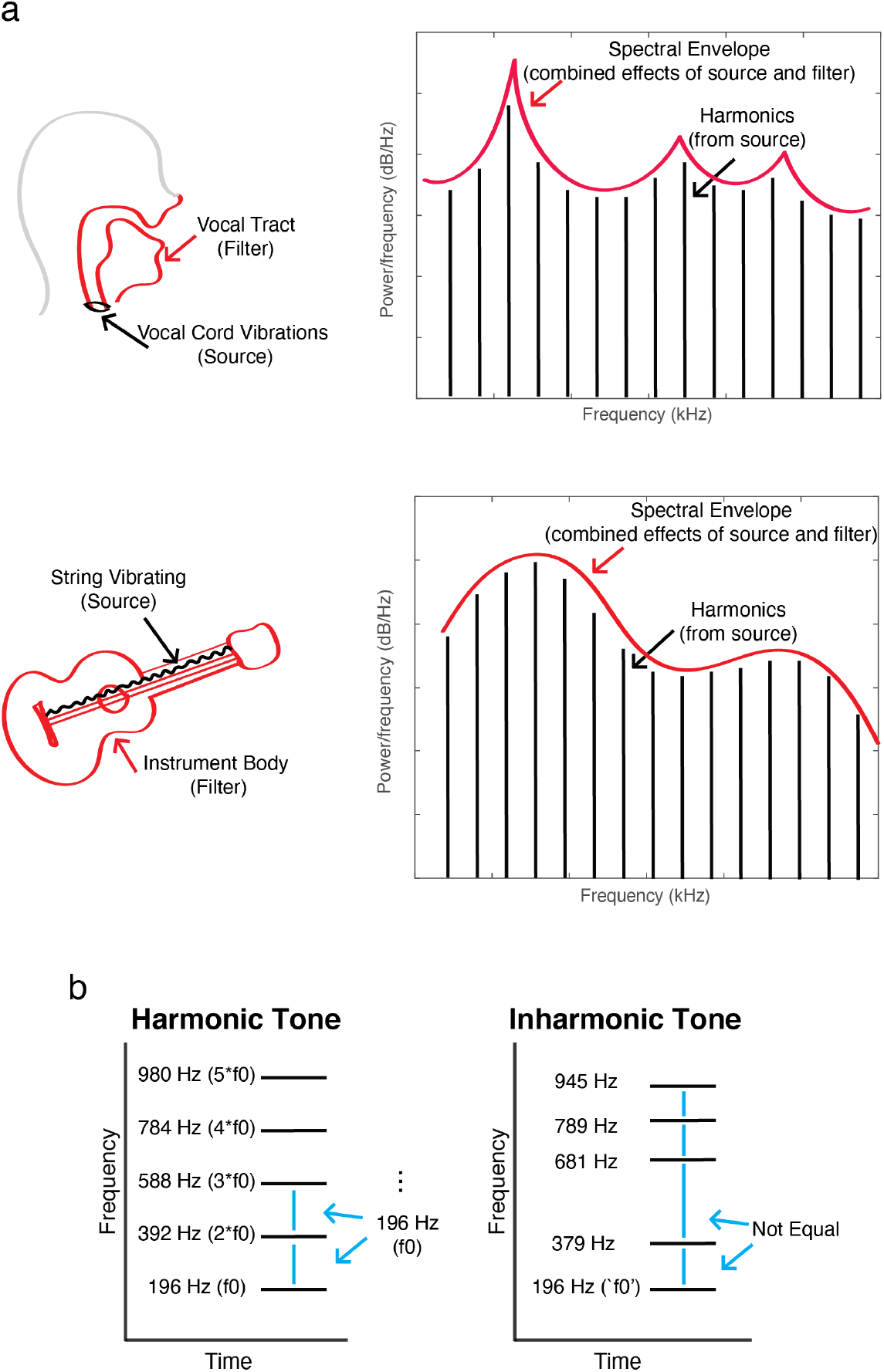
Acoustic variables related to pitch perception. a. Schematic demonstrating the source-filter generative model of many speech and instrument sounds. Vibrations of the vocal cord or a string on a guitar, for example, produce harmonics which are then filtered by the vocal tract or instrument body, respectively. Filtering changes the overall shape of the power spectra, i.e. the spectral envelope. As a result, sounds can have the same fundamental frequency (f0), but different spectral envelopes. b. Schematic spectrogram of a 196 Hz Harmonic tone (G3 in the Western classical scale), and a related inharmonic tone. The inharmonic tone was generated by altering the frequencies of all harmonics above the f0, by sampling a jitter value from the distribution U(−0.5, 0.5), multiplying that jitter by the f0, then adding the resulting value to the frequency of the respective harmonic, constraining adjacent harmonics to be separated by at least 30 Hz (via rejection sampling) in order to avoid salient beating. For the purposes of this paper, such an inharmonic tone will be described as having a “nominal f0” of 196 Hz, as it is generated by perturbing the harmonics of 196 Hz.

One example of this invariance is the ‘missing fundamental illusion’. When the lowest frequency partial is removed from a harmonic tone, the f0 of the tone remains the same, and listeners are thought to be able to hear this invariant property of the tones even though the spectral envelope has changed (Licklider, 1954). Many studies have explored variants of this phenomenon with synthetic tones, testing pitch matching or discrimination of tones that vary in their spectra. Human pitch discrimination exhibits some robustness to spectral envelope variation, in that listeners are above chance at discriminating tones that have non-overlapping sets of harmonics (Micheyl & Oxenham, 2004; Moore et al., 1992; Singh & Hirsh, 1992). However, discrimination is nonetheless impaired compared to when tones have the same set of harmonics (Allen & Oxenham, 2014; Melara & Marks, 1990; Micheyl & Oxenham, 2004; Moore et al., 1992; Moore & Glasberg, 1990; Russo & Thompson, 2005; Singh & Hirsh, 1992; Warrier & Zatorre, 2002). Similar effects appear to occur with natural spectral envelope variation, as is present between different syllables or instruments. Two previous studies compared pitch judgments of pairs of notes played on same vs. different instruments (Vurma et al., 2011; Zarate et al., 2013), finding slight deficits in performance when notes were from different instruments. Several previous experiments have likewise examined discrimination of pitch among different vowels, finding deficits in pitch discrimination between different vs. the same vowels (Chuang & Wang, 1978; Hellström et al., 1994; Kuhl & Miller, 1982; Miller, 1978; Repp & Lin, 1990; Stoll, 1984). At face value these findings are at odds with the idea that pitch is invariant to the spectral envelope. But these studies left open the basis of the observed deficits, as well as the basis of the residual invariance.

The fact that relative pitch judgments can be made in two different ways (using either frequency partials or f0), coupled with the common assumption that f0 representations are invariant to the spectral envelope, raises the question of whether any invariance of relative pitch would depend on whether listeners rely on f0 representations or not. The purpose of this paper was to investigate this question. One tool to test for representations of the f0 is to compare performance with harmonic sounds to that with inharmonic sounds, which lack a single f0 in the range of audible pitch (Figure 1b). A role for f0 representations in spectral invariance has been suggested by discrimination advantages for harmonic tones over inharmonic tones when the tones being compared contain distinct sets of harmonics (Micheyl et al., 2010; Moore & Glasberg, 1990). However, these studies used extreme degrees of spectral envelope variation, unlike that which occurs for natural sounds. It was a priori unclear whether this harmonic advantage would be observed for naturally occurring spectral envelope variation, as when comparing the pitch of different vowels in a speech utterance, or the pitch of different instruments.

The specific goals of this paper were to assess 1) whether any invariance of pitch perception to naturally occurring timbre differences is dependent on representations of f0 and 2) to what extent any limits on invariance reflect imperfect invariance of pitch representations themselves vs. interference in the judgments that operate on those representations.

To examine the first question, we measured f0 discrimination with both harmonic and inharmonic versions of the same or different spoken vowels, as well as with notes played on the same or different musical instruments. Our assumption was that if a task relies on representations of the f0, performance should be impaired with inharmonic stimuli (Faulkner, 1985; McPherson et al., 2022; McPherson & McDermott, 2018; McPherson & McDermott, 2020; Micheyl et al., 2010; Moore & Glasberg, 1990). Specifically, if spectral invariance is mediated by representations of an estimate of the f0 (henceforth referred to as “representations of f0”, that mediate “f0-based pitch”), we predicted that there would be harmonic advantages when discriminating sounds that differ in their spectral envelopes.

To answer the second question, we measured pitch discrimination across short time delays, leveraging the apparent robustness of f0 representations over time (McPherson & McDermott, 2020). Our hypothesis was that if limits to spectral invariance derive from bias in the auditory system’s representation of f0, such bias should remain evident in discrimination across a delay. If instead the limits to invariance arise only at a subsequent comparison stage (e.g. if independent representations of the spectral envelope and the f0 interfere when making comparisons), effects of the spectral envelope on discrimination might decrease across a delay, as the representation of the spectral envelope might degrade more rapidly than that of the f0.

We found that discrimination was only partially invariant to real-world differences in spectral envelope between different vowels or instruments. Judgments were biased depending on whether the spectral envelope shifted congruently or incongruently with the f0, consistent with previous studies (Allen & Oxenham, 2014; Chuang & Wang, 1978; Hellström et al., 1994; Kuhl & Miller, 1982; Micheyl & Oxenham, 2004; Miller, 1978; Moore & Glasberg, 1990; Repp & Lin, 1990; Siedenburg et al., 2022; Singh & Hirsh, 1992; Stoll, 1984). However, invariance was similarly limited for harmonic and inharmonic tones, indicating that invariance does not depend on representations of f0 (addressing the first question). This bias decreased when we introduced delays between harmonic tones, suggesting that representations of the f0 and spectral envelope decay at different rates, and are thus independent, and that the observed biases occur at a comparison stage that cannot separate the effects of pitch and timbre (addressing the second question). Overall, the results suggest that representations of the f0 are relatively invariant to spectral envelope. But relative pitch judgments are not, even when the variation in spectral envelope is naturally occurring, and even when such judgments are reliant on representations of the f0.

## 2 METHODS AND MATERIALS

This section contains aspects of the methods that were shared by two or more experiments.

### 2.1 Ethics

All experiments were approved by the Committee on the use of Humans as Experimental Subjects at the Massachusetts Institute of Technology and were conducted with the informed consent of the participants.

### 2.2 Audio presentation and procedure for online experiments (Experiments 1-2, 4-5)

Experiments 1, 2, 4, and 5 were completed online due to the COVID-19 pandemic. Online psychoacoustic experiments sacrifice control over absolute sound presentation levels and spectra, but we have repeatedly found that online experiments replicate results obtained in controlled laboratory conditions provided that modest steps are taken to help ensure reasonable sound quality and compliance with task instructions (Kell et al., 2018; McPherson et al., 2020; McPherson et al., 2022; McPherson & McDermott, 2020; McWalter & McDermott, 2019; Traer et al., 2021; Woods & McDermott, 2018). Experiments were conducted on the Amazon Mechanical Turk platform. We limited participation to individuals with US-based IP addresses. Before beginning the experiment, potential participants gave consent to participate and were instructed to wear headphones and ensure they were in a quiet location. They then used a calibration sound (1.5 seconds of speech-shaped noise) to set their audio presentation volume to a comfortable level. The experimental stimuli were normalized to 6 dB below the level of the calibration sound to ensure that they were likely to be audible without being uncomfortably loud. Participants then completed a brief screening test designed to help ensure they were wearing earphones or headphones (Woods et al., 2017). If they passed this screening, they could continue to the main experiment. To incentivize good performance, online participants received a compensation bonus proportional to the number of correct trials. All online experiments began with a set of screening questions that included a question asking the participant if they had any hearing loss. Anyone who indicated any known hearing loss was excluded from the study. Across all the online experiments in this paper, 3.5% of the 1052 participants who initially enrolled self-reported hearing loss and were excluded (all but four of these individuals also failed the headphone screening, and would have been excluded regardless). All participants whose data were analyzed thus self-reported normal hearing.

For technical reasons all stimuli for online experiments were generated ahead of time and were stored as .wav files on a university server, from which they could be loaded during the experiments. 20 sets of stimuli were pre-generated for each experiment, and participants only heard stimuli from one of these sets, randomly assigned. Sets differed based on the specific f0s used in each trial (randomly selected based on parameters described for each experiment), the specific jitters used to make sounds inharmonic (in experiments where we tested inharmonic tones), and the specific vowel (Experiment 1), or Instrument (Experiment 2) exemplars.

### 2.3 Inharmonic stimuli

In order to make synthetic tones, spoken syllables, or musical instrument notes inharmonic, the frequency of each harmonic, excluding the fundamental, was perturbed (jittered). For syllables and instrument notes, the original harmonic frequencies were derived from the f0 estimated from the original recording. In all experiments, frequencies were jittered by an amount chosen randomly from a uniform distribution, *U*(−.5, .5). These jitter values were multiplied by the f0 of the tone and added to the frequency of the respective harmonic. For example, if the f0 was 200 Hz and a jitter value of −0.39 was selected for the second harmonic; its frequency would be set to 322 Hz (200*2+200*-.39, Figure 1b). To minimize salient differences in beating, jitter values were constrained via rejection sampling such that adjacent harmonics were always separated by at least 30 Hz. Jitter values were generated for each harmonic in succession, beginning with the second, subject to the 30 Hz constraint. A different jitter pattern was chosen randomly for each trial, but the same jitter pattern was applied to the two tones/notes/vowels within a trial.

### 2.4 Distortion products

Distortion products can in principle explain differences in performance for harmonic and inharmonic stimuli (because they should be stronger in harmonic stimuli). However, in most cases in which we compared task performance for harmonic and inharmonic stimuli (Experiments 1, 2, and 5), the stimuli contained all lower harmonics, including the fundamental. Because distortion products are typically substantially lower in level than the stimulus components that generate them (Norman-Haignere & McDermott, 2016; Pressnitzer & Patterson, 2001), they are unlikely to be detectable in stimuli that contain all lower harmonics, and thus are unlikely to account for any differences between performance for harmonic and inharmonic stimuli. We did not include masking noise in the stimuli for those three experiments. The exception was Experiment 3, where some of the stimuli were missing the lower harmonics. In this experiment we included masking noise to rule out distortion products as the basis for discrimination.

### 2.5 Feedback

Participants were asked to report whether the second note in a trial was ‘higher’ or ‘lower’ than the first, and we did not specify that listeners should explicitly ignore timbre. However, we provided feedback corresponding to changes in f0 (or nominal f0, in the inharmonic cases) for all tasks except Experiment 3 (where we anticipated chance or below-chance performance for most of the Inharmonic conditions, and we did not want participants to get discouraged).

### 2.6 Statistics

Data distributions were evaluated for normality by visual inspection and parametric statistics were used across all conditions. Unless otherwise noted we tested hypotheses using repeated measures ANOVAs. All analysis was completed in MATLAB (version 2020b). Bayesian statistics were used to evaluate null results (*JASP*, 2020).

## 3 RESULTS

### 3.1 Experiment 1: Pitch discrimination with same vs. different vowels

#### 3.1.1 Purpose and procedure

We began by testing up/down discrimination of same vs. different vowels. During each trial, participants heard two vowels and judged whether the second was higher or lower in pitch than the first, indicating their response by clicking one of two buttons (‘higher’ or ‘lower’, Figure 2a). The conditions resulted from fully crossing three variables: Same vs. Different vowels, Harmonic vs. Inharmonic, and 4 frequency differences (.33, 1, 3, and 9 semitones). Conditions were randomly intermixed during the experiment, and participants completed 30 trials per condition.

**Figure 2:**
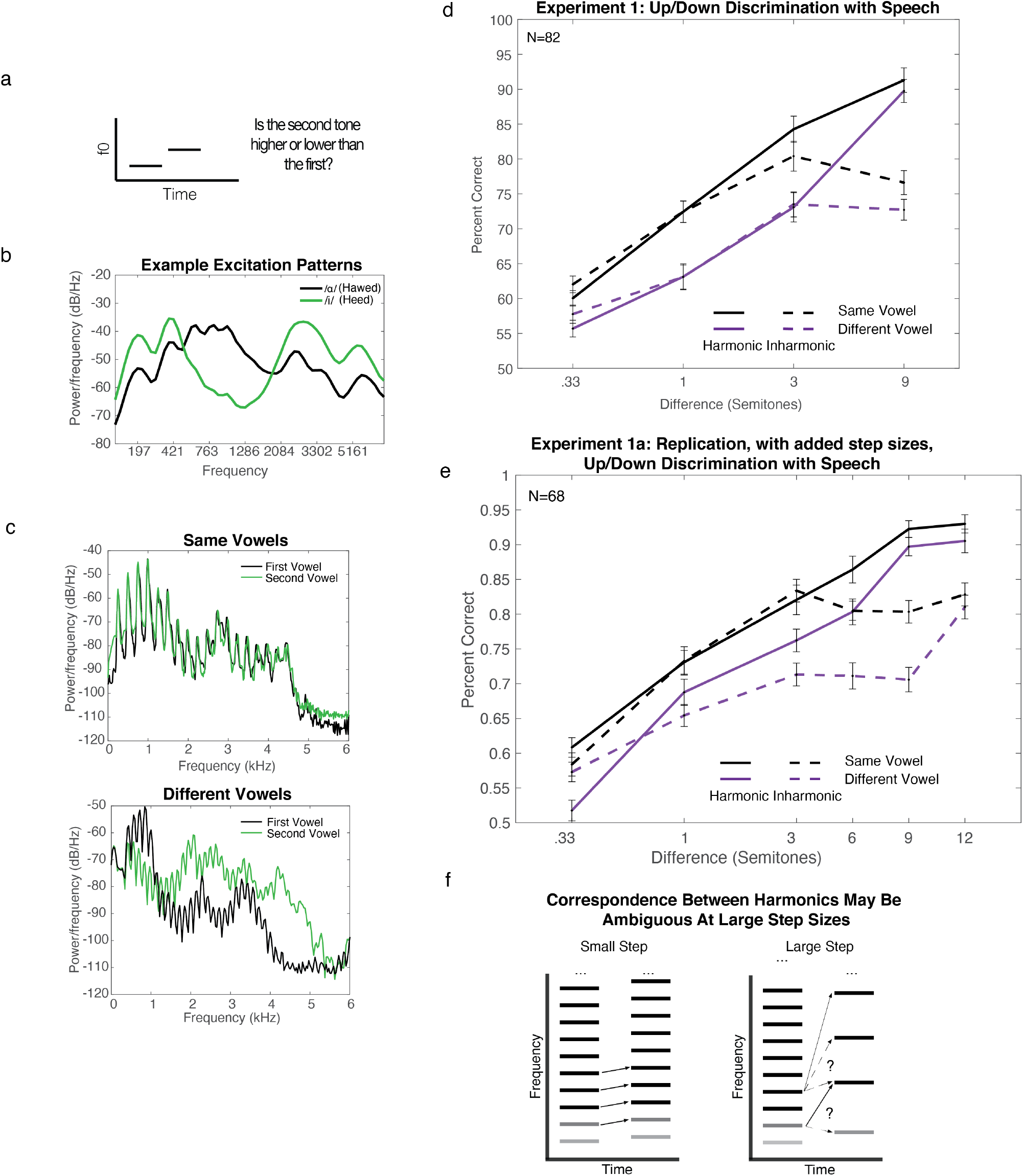
Stimuli and Results for Experiment 1, Pitch discrimination with same vs. different vowels. a. Task for all experiments. Participants heard two sounds and judged which was higher in pitch. b. Example excitation patterns for two vowels (x-axis scaled in Equivalent Rectangular Bandwidths based on Glasberg and Moore (1990)). Excitation patterns contain peaks for individual harmonics, but also reveal the differences in spectral envelope between the vowels c. Example stimulus power spectra from two exemplars of the same vowel (top), and two different vowels (bottom). d. Results from Experiment 1a. Here and in e, error bars denote standard error of the mean. e. Results from Experiment 1b. f. Schematic illustrating potential difficulty in matching harmonics between tones with a fixed filter that are separated by a large step size. Gray level of line segments denotes amplitude (lower harmonics are attenuated by the filter). To facilitate inspection of the ambiguity in matching, only lower portion of spectrum is shown. When two tones have similar f0s, the correspondence between harmonics is relatively unambiguous (left), but when the f0 change is large enough the correspondence between harmonics is ambiguous (right), potentially impairing performance in conditions in which listeners base judgments on shifts in harmonics rather than the f0.

Our hypothesis was that any spectral invariance (ability to discriminate pitch despite differences in the spectra) would be mediated by representations of the f0, which would be indicated by a harmonic advantage specific to (or greater when) discriminating different vowels.

#### 3.1.2 Stimuli

During each trial, participants were asked to compare two vowels, each spoken by the same speaker to maximize ecological validity (we envisioned the task as potentially tapping into the mechanisms underlying the perception of the prosodic contour of an utterance). We aimed to choose a set of pairings for which the two vowels in each pair varied substantially in their spectral envelope (Figure 2b-c; example same vs. different vowel spectra).

To find vowel pairs that differed in their spectral envelope, we compared the excitation patterns across pairs of different vowels with the same f0. Excitation patterns were calculated by passing waveforms (each normalized to the same RMS level) through a gammatone filter bank (Slaney, 1998) approximating the frequency selectivity of the cochlea. The filter bank was implemented with Malcolm Slaney’s Auditory Toolbox, with the lowest center frequency set to 50 Hz, the number of channels set to 64, and the highest center frequency set to 7576 Hz, with filters evenly spaced on an equivalent-rectangular-bandwidth scale (Glasberg & Moore, 1990). The excitation pattern was calculated from a magnitude spectrogram generated from these filter outputs (the RMS amplitude within .025 s bins; hop size of .01 s) by averaging the spectrogram over time. The spectral envelope difference between two vowels was calculated as the sum over frequency of the absolute value of the dB differences between the two excitation patterns (Figure 2b).

Vowels were selected from the Hillenbrand vowel set (Hillenbrand et al., 1995), which contains recordings from 45 men, 48 women, and 46 children (the latter group was not used in this experiment) each producing 12 vowels in h-V-d syllables. We included the vowels /I, ɪ, æ, ɑ ɔ, ε, u, ʌ, ʊ, ɝ/ in our analysis, omitting the two diphthongs in the Hillenbrand set (/ou/ and /eɪ/) because their spectra are not fixed across their duration. We calculated the spectral envelope difference between every pair of the vowels (45 pairs) for every speaker (all 45 male speakers and 48 female speakers). Before comparing the spectra of each vowel in a pair, we pitch-shifted the two vowels to match their mean f0 (we pitch-shifted each of the two vowels by and equal and opposite amount, using STRAIGHT, described below). We settled on the set of 10 vowel pairings that maximized the average spectral envelope difference between vowels of the same speaker. We did not otherwise constrain the vowel pairings, so some vowel tokens are over-represented (see Supplementary Table 1 for the final set of vowels).

For every ‘Different Vowel’ condition, we used each of the 10 vowel pairings 3 times, randomly sampling from the speakers in the set. 15 of these 30 vowel sets were selected from female speakers, and 15 from male speakers. The presentation order of the two vowels within a trial was randomized.

For ‘Same Vowel’ conditions, we chose a single vowel recording to use for both intervals of the trial. We sampled a set of 30 vowels that included all 20 of the vowels from the 10-vowel pairings set along with an additional 10 vowels that were sampled randomly from the set of 20 without replacement. We again sampled randomly from the speakers in the set, balancing for speaker gender.

To make the final stimuli, vowels were pitch-shifted and resynthesized to be either harmonic or inharmonic using the STRAIGHT analysis and synthesis method (Kawahara, 2006; Kawahara & Morise, 2011; McDermott et al., 2012). STRAIGHT decomposes a recording of speech or instruments into periodic and aperiodic excitation and a time-varying filter. If the periodic excitation is modeled as a sum of sinusoids, one can alter the frequencies of individual harmonics, and then recombine them with the unaltered aperiodic excitation and filter to generate harmonic or inharmonic speech. This manipulation leaves the spectral envelope of the speech largely intact, and a previous study suggests that intelligibility of inharmonic speech in quiet is comparable to that of harmonic speech (Popham et al., 2018).

We pitch-shifted vowels to achieve the desired f0 difference, but did not otherwise modify the pitch contours of the vowels. The natural vowels had pitches ranging from 59.7 Hz to 270.0 Hz. The pitches were bimodally distributed (due to the sex of the speakers), with the mean f0 for male speakers occurring at 132.6 Hz and the mean for female speakers occurring at 220.2 Hz. The pitch contours of the vowels fluctuated with an average standard deviation of .18 of a semitone. To perform the pitch adjustments, we first calculated the combined mean f0 across both vowels. We pitch-shifted the vowels so that the mean f0 difference between them was .33, 1, 3, 6, 9 or 12 semitones, depending on the trial, such that the final signals were equidistant from the original combined mean f0 (i.e. we preserved the mean f0 of the two vowels). This ensured that the pitch of both vowels was shifted as little as possible from the original pitch, and by equal amounts.

Because we did not pitch-flatten or otherwise modify the pitch contours of the vowels, it seemed important to control for the fact that in the Different Vowel case, the two selected vowels would have different pitch fluctuations. We addressed this possible confound by giving the two intervals in a Same Vowel trial different pitch contours. This was achieved by replacing the pitch contour of one of the two vowels in the Same Vowel condition with the pitch contour from what would have been the other vowel, were it a Different Vowel condition. For example, if the vowel in a Same Vowel trial was the vowel /æ/ drawn from the pair /æ/ and /u/, we would use a single recording of the /æ/ vowel from one speaker for both intervals in the trial, but would replace the f0 contour for one of the intervals with that from the speaker’s /u/ vowel. In this way, the pitch variability within trials was matched between Same Vowel and Different Vowel conditions.

The resynthesized vowels were truncated at 400 ms, with onsets and offsets shaped by a 15 ms half-Hanning window, and presented in sequence (with no time delay between vowels). Sounds were sampled at 16 kHz (the sampling rate of the Hillenbrand vowel recordings).

Stimuli for Experiment 1b were identical to those in Experiment 1a, with the addition of a 6 and 12 semitone condition.

#### 3.1.3 Participants

127 participants passed the headphone check and enrolled in Experiment 1a online. 45 had overall performance (averaged across all conditions) below 55% correct, and were removed from further analysis, leaving 82 participants whose data are reported here. 26 identified as female, 56 as male, 0 as non-binary. The average age of these participants was 38.0 years (S.D.=10.4). 34 participants had four or more years of musical training (self-reported), with an average of 11.4 years (S.D.=8.4).

We based our initial target sample size on the same pilot experiment and power analysis used for Experiment 2 (described below). This pilot experiment was conducted in-lab, and listeners were asked to discriminate tones played on the same or different instruments. It showed that we needed 7 participants to have a 95% chance of observing an effect of same vs. different instruments at a significance level of .01. As we observed a null effect of harmonicity for small f0 differences after collecting our initial data, we continued data collection until Bayesian statistics converged on support for or against the null hypothesis. Unlike frequentists statistics, Bayesian statistics can converge on supporting the null hypothesis as additional data is added to the analysis (Rouder et al., 2009).

118 participants passed the headphone check and enrolled in Experiment 1b online. 50 participants had overall performance (averaged across all conditions) below 55% correct, and were removed from further analysis, leaving 68 participants. 19 identified as female, 48 as male, 1 as non-binary. The average age of the participants was 37.8 years (S.D.=10.5). 19 participants self-reported four or more years of musical training, with an average of 8.9 years (S.D.=6.0).

The sample size for Experiment 1b was based on pilot data from 36 participants. The pilot experiment contained only step sizes 3, 6, 9, and 12 semitones. Based on this pilot data, we aimed to collect data from 68 participants to have a 95% chance of seeing a significant difference between Same and Different Vowels for the Harmonic stimuli at a 6-semitone step size, at a significance level of *p*<.01.

#### 3.1.4 Experiment 1a: Results and discussion

Our first hypothesis was that participants are invariant to spectral envelope differences between vowels, such that we would observe similar discrimination across same and different vowels. Instead, we found that discrimination was worse when participants compared different vowels than when they compared instances of the same vowel (main effect of Same vs. Different vowels, F(1,81)=78.76, *p*<.0001, η_p_^2^=.49, Figure 2d). However, participants were nonetheless well above chance when discriminating the pitch of different vowels. Listeners thus exhibited some degree of spectral invariance.

Our second hypothesis was that harmonicity would be critical to any observed spectral invariance, such that there might be a harmonic advantage in the different vowel condition but not the same vowel condition. Contrary to this hypothesis, there was no interaction between the effects of Same vs. Different vowels and Harmonicity (F(1,81)=0.20, *p*=.66, η_p_^2^=.002, the BF_incl_=0.76, specifying a multivariate Cauchy prior on the effects, provided anecdotal support for the null hypothesis (*JASP*, 2020; Wagenmakers et al., 2018)). In particular, for the three smallest step sizes, there was no difference between Harmonic and Inharmonic performance for either the Same Vowel or Different Vowel conditions (no main effect in either case; Same Vowel: F(1,81)=0.37, *p*=.55, η_p_^2^=.005, Different Vowel: F(1,81)=0.66, *p*=.42, η_p_^2^=.008). The Bayes factors in each case (BF_incl_ of .13 and .11 for Same and Different Vowels, respectively) provided moderate support for the null hypotheses (*JASP*, 2020; Wagenmakers et al., 2018).

However, there were clear differences between performance with Harmonic and Inharmonic sounds for both vowel conditions when the step size was large. At a 9 semitone difference there was a highly significant effect of Harmonicity (F(1,81)=104.91, *p*<.0001, η_p_^2^=.56; this drove an overall effect of Harmonicity across all step sizes (i.e. the f0 difference, or nominal f0 difference in the inharmonic case, between the two sounds on a trial, in semitones), F(1,81)=18.94, *p*<.0001, η_p_^2^=.19). This effect with large intervals resulted in a highly significant interaction between step size and Harmonicity (F(3,243)=50.71, *p*<.0001, η_p_^2^=.38). This result is similar to one that we previously found for synthetic tones with a fixed filter (replotted for convenience in Supplementary Figure 1), and suggests the effect has relevance for real-world listening conditions. The results with larger intervals are further explored in Experiment 1b, described below (Figure 2e).

All together, these results suggest that participants are not fully invariant to changes in timbre, and that the invariance they have is not mediated by a representation of f0. However, representations of f0 appear to be helpful in discriminating large pitch differences. For large pitch differences it may be difficult for listeners to match the frequency partials of one inharmonic tone to those of the next inharmonic tone, as is necessary to determine the step from a representation of differences between individual partials (Figure 2f). Having access to the f0 (as is available in harmonic tones) may help listeners resolve these ambiguities.

#### 3.1.5 Experiment 1b: Replication with additional pitch differences

One concern regarding the results in Experiment 1a was that at the 9-semitone pitch difference for which there was a clear advantage for the Harmonic over the Inharmonic condition, there was only a small (and statistically insignificant) difference between the Same and Different vowel conditions for harmonic stimuli (t(81)=0.84, *p*=0.40). Although a difference between Same and Different vowel conditions was evident at smaller intervals, these intervals produced no difference between Harmonic and Inharmonic conditions. The results thus did not provide evidence that judgments based on f0 representations are impaired by differences in the spectral envelope. However, it seemed possible that performance in the 9-semitone condition was influenced by ceiling effects, such that a somewhat smaller interval might show a Harmonic advantage while also showing a Same vowel advantage, which would provide evidence that both regimes of relative pitch judgments are impaired by spectral envelope variation. We therefore replicated Experiment 1a, but added two additional step sizes, 6 and 12 semitones. The 12-semitone condition was included to test whether listeners were at ceiling in the 9-semitone condition, in which case performance should be the same when we increased the size of the interval further.

At six semitones, we observed a significant advantage for Harmonic vs. Inharmonic stimuli (t(67)= 5.49, *p*<.0001), as well as a significant advantage for Same vs. Different Vowels when sounds were harmonic (t(67)=4.38, *p*<.0001). This suggests that even when listeners are relying on f0-based pitch (evidenced by the harmonic advantage), they are not fully invariant to the spectral envelope variations in vowel identity (evidenced by the advantage for Same vs. Different Vowels). There was no significant difference between Harmonic stimuli at 9 and 12 semitones (no main effect of step size, F(1,67)=0.46, *p*=.50, η_p_^2^=.007), suggesting listeners are indeed at ceiling in both conditions. This ceiling effect likely explains why we did not observe the effect of Same vs. Different Vowels at the 9-semitone step size in Experiment 1a.

### 3.2 Experiment 2: Pitch discrimination with notes played on the same vs. different instruments

#### 3.2.1 Purpose and procedure

The procedure and hypothesis for Experiment 2 were analogous to those of Experiment 1. However, instead of hearing pairs of vowels, participants heard pairs of instrument notes that were resynthesized to be either harmonic or inharmonic. Participants again completed 30 trials per condition.

#### 3.2.2 Stimuli

Each trial contained two notes resynthesized from recordings of instruments. To approximately replicate the maximal spectral envelope difference between instruments that a listener might hear in Western music, we analyzed the spectra of different instruments to identify a subset that differed the most in their spectra, using the same excitation pattern analysis procedure used with vowels in Experiment 1. We then selected pairs of instruments from this subset for the Different Instrument trials, and single instruments from this subset for Same Instrument trials.

Recordings were drawn from the RWC Music Database of Musical Instrument Sounds (Goto et al., 2003). We excluded instruments whose recordings did not include all notes in the Western Scale within the approximately 200-400 Hz range we were planning to test in the experiment. This criterion excluded several non-Western instruments (Sho, Koto, etc.) and instruments with low or high pitch ranges (contrabass, soprano recorders etc.). Percussion instruments were also excluded because they are naturally inharmonic. These exclusions left us with 25 instruments (see Supplementary Table 2). We then searched through all instrument pairings to find a set of instruments that maximized variability in timbre.

For each pairing, we compared the excitation patterns of notes with the same f0, for all notes between G3 and G4 (196-392 Hz). Sets of 5 instruments were then ranked by the pairwise spectral envelope difference summed across all pairs of instruments within the set, and the combination that maximized this quantity was selected. We settled on an instrument set size of 5 because increasing the set size above 5 decreased the average variability between instruments. Because plucked instruments are not perfectly harmonic, it seemed wise not to have too many of them (even though all sounds were resynthesized to be harmonic for the experiment), so we constrained the sets of instruments so that they could only contain one plucked instrument. The final set included baritone saxophone, cello, oboe, pipe organ, and ukulele (Figure 3a). There were 10 possible pairs of instruments that could be drawn from this set. We adopted this procedure (in contrast to that used in Experiment 1, where we selected 10 pairs of vowels with substantial spectral envelope differences) because the spectral envelope differences between instruments were sufficiently pronounced that it was possible to find a set of 5 for which all pairings were between instruments with distinct timbres.

**Figure 3:**
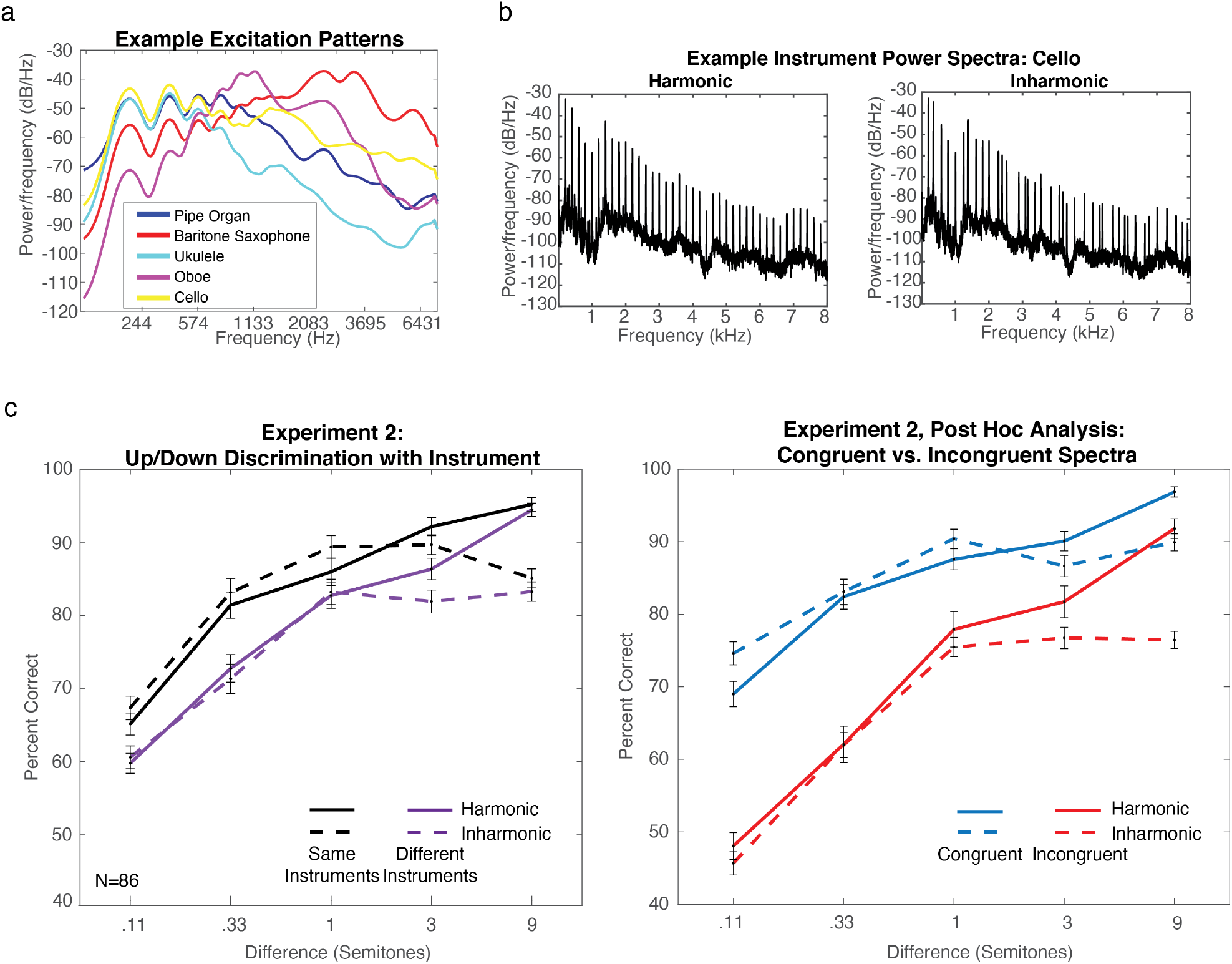
Stimuli and results for Experiment 2, Pitch discrimination with notes played on the same vs. different instruments. a. Example excitation patterns for notes with the same f0 (200 Hz) played on the five instruments used in Experiment 2. Excitation patterns contain peaks for individual harmonics, but also reveal the differences in spectral envelope between the instruments. b. Example power spectra of harmonic and inharmonic instrument notes used in Experiment 2. Note the irregularly spaced frequencies in the inharmonic notes, due to the frequency jittering. c. Results from Experiment 2. Error bars denote standard error of the mean. Left panel shows Same and Different Instrument (main results). Right panel shows the results of a post-hoc analysis in which ‘Different Instrument’ trials were grouped by whether the spectral centroid of the notes moved congruently or incongruently with the f0 shift (‘Congruent’ and ‘Incongruent’ subsets of trials).

To generate individual trials, the first note was randomly selected from a uniform distribution over the notes in a Western classical chromatic scale between G3 and G4 (196-392 Hz). A recording of this note, from a randomly selected instrument, was chosen as the source for the first note in the trial. The second note was drawn from either the same instrument or a different instrument, depending on the condition. If different, the second instrument was drawn at random from the remaining 4 instruments. The recording whose f0 was closest to that needed for the intended frequency/f0 difference was chosen to generate the second stimulus. For differences less than one semitone, a note one semitone above or below the first note was chosen as the source for the second note in the trial. We tested five different step sizes: .11, .33, 1, 3 and 9 semitones. We included one step that was smaller than those in Experiment 1 because we pitch flattened the instrument stimuli, making the task easier.

The two notes were then modified using the STRAIGHT analysis and synthesis method (Kawahara, 2006; Kawahara & Morise, 2011; McDermott et al., 2012), in the same way we modified vowels in Experiment 1, to conform to the stimulus parameters for the condition. We confirmed by eye that the spectral envelopes estimated by STRAIGHT for the instruments in the set were reasonable in all cases. The same spectral envelopes were used for harmonic and inharmonic notes, such that any deviations from the true spectral envelopes of the instruments would not influence the results. After analysis, the f0 contours were flattened to remove any vibrato and shifted to ensure that the f0 differences were exact. The notes were then resynthesized with harmonic or inharmonic excitation (Figure 3b). We shifted both notes by the same amount to achieve the desired f0 differences (in other words, when the f0 difference of the source notes was less than the f0 difference for the condition, the lower note’s f0 was decreased and the higher note’s f0 was increased, and vice versa). As a result of the resynthesis, the small deviations from perfect harmonicity found in natural instruments were removed for the harmonic conditions (Fletcher, 1999). The resynthesized notes were truncated at 400 ms, windowed with a 15 ms half-Hanning window, and presented sequentially with no delay between notes. Sounds were sampled at 16 kHz.

#### 3.2.3 Participants

105 participants passed our headphone check and completed Experiment 2 online. 19 had overall performance (averaged across all conditions) below 55% correct and were removed from further analysis. Of the 86 remaining participants, 25 identified as female, 61 as male, 0 as non-binary. The average age of these participants was 37.4 years (S.D.=10.7). 32 participants had four or more years of musical training (self-reported), with an average of 10.6 years (S.D.=10.0).

The sample size was ultimately determined by the desire to provide evidence for or against the null hypothesis that discrimination was the same for harmonic and inharmonic stimuli. We initially performed a power analysis with pilot data. The pilot experiment had slightly different stimuli (same sets of instruments, but instrument notes were high-passed filtered and had low-pass noise added to mask the f0 component), did not include a 9-semitone step size, and was run in the lab. This pilot experiment showed a main effect of same vs. different instruments (η_p_^2^=.64), but no effect of harmonicity. We thus wanted to be well-powered to test for an effect of same vs. different instrument, but also to provide evidence for a null effect of harmonicity should it hold. The power analysis (using G*Power) indicated that we would need only 7 participants to observe the effect of same vs. different instruments at a significance level of .01, 95% of the time (Faul et al., 2007). We aimed to have at least this many musicians (with four or more years of musical training) and non-musicians (fewer than four years of musical training), to be able to analyze the groups separately (see *Effects of Musicianship* section below). After replicating the null effect of harmonicity for small f0 differences in the first 7 participants who competed the experiment, we then continued data collection (in sets of approximately 8-12 participants) until Bayesian statistics converged on support for or against the null hypothesis.

#### 3.2.4 Results and discussion

As in Experiment 1, our first hypothesis was that participants might be invariant to differences between instruments. Contrary to the predictions of strict invariance, and analogous to the result of Experiment 1, we found that discrimination was worse when participants compared notes from different instruments than when they compared notes from the same instruments (main effect of same vs. different instruments, F(1,85)=140.34, *p*<.0001, η_p_^2^=.62, Figure 3c, Same vs. Different instruments plotted on separate graphs to ease comparison for a subsequent analysis, described below). Participants were nonetheless well above chance at discriminating notes from different instruments (performance was above chance even for the smallest pitch change tested, .11 semitones, t-test against .5, t(85)=10.04, *p*<.0001), again demonstrating some degree of spectral invariance. Overall, thresholds were lower in Experiment 2 compared to Experiment 1, as expected given that the stimuli in Experiment 2 were pitch-flattened, whereas vowels in Experiment 1 retained their natural pitch fluctuations.

Our second hypothesis was that harmonicity would be critical to spectral invariance, such that there would be a harmonic advantage in the different instrument condition but not in the same instrument condition (i.e., an interaction). There was a statistically significant interaction between Harmonicity and Same vs. Different instruments (F(1,85)=11.60, *p*=.001, η_p_^2^=.12). However, the effect was small, and was mainly driven by better performance for Inharmonic notes in the Same Instrument condition (there was a significant interaction for the three smallest semitones, F(1,85)=7.50, *p*=.007, η_p_^2^=.08, but not for the largest two, F(1,85)=3.68, *p*=.06, η_p_^2^=.04). As with vowels in Experiment 1, there was no significant effect of Harmonicity with Different Instruments for the smallest three step sizes (F(1,85)=0.002, *p*=.96, η_p_^2-^ <.0001), and evidence for the null hypothesis (Bayes factor BF_incl_, specifying a multivariate Cauchy prior on the effects, was .07, providing strong support of the null hypothesis (*JASP*, 2020; Wagenmakers et al., 2018)), indicating that there was no more invariance to the spectral envelope when listeners could have used representations of the f0. There was a significant effect of Harmonicity for Same Instruments with small step sizes (F(1,85)=15.66, *p*=.0002, η_p_^2^=.16), but it was due to performance being marginally better for Inharmonic notes, for reasons that are unclear. And like Experiment 1b, for the moderate step size (3 semitones), there was both an advantage for Harmonic stimuli over Inharmonic stimuli (t(85)=5.16, *p*<.0001, averaged across Same and Different instrument conditions), and an advantage for Same vs. Different stimuli in the Harmonic condition (t(85)=5.41, *p*<.0001), indicating incomplete invariance even when listeners were likely relying on representations of f0. Overall, the results are again inconsistent with the hypothesis that representations of f0 support spectral invariance for natural sounds.

As with vowels in Experiment 1, discrimination was worse for Inharmonic notes when the step size was large. At a 9 semitone difference there was a highly significant effect of Harmonicity (F(1,85)=136.71, *p*<.0001, η_p_^2^=.62) This difference drove an overall effect of Harmonicity across all step sizes, F(1,85)=22.16, *p*<.0001, η_p_^2^=.21), but there was again a highly significant interaction between step size and Harmonicity (F(1,85)=48.93, *p*<.0001, η_p_^2^=.37).

All together, these results suggest that pitch discrimination is not completely invariant to naturally occurring differences in spectral envelope, and that the partial invariance of pitch discrimination is not mediated by representations of f0. Consistent with prior work, it appears that listeners use shifts in individual partials to register small pitch changes (yielding the matched performance for Harmonic and Inharmonic conditions). But they transition to using the f0 for large changes (better performance for Harmonic conditions, irrespective of Same vs. Different spectral envelope).

#### 3.2.5 Post Hoc Spectral Centroid Analysis

It seemed plausible that the deficit we observed in the Different Instrument condition could depend on whether the difference in the spectral center-of-mass (centroid) between the two instrument notes was congruent or incongruent with the pitch change. To investigate this issue, we performed an additional analysis in which we calculated the spectral centroid of each note and then separated Different Instrument trials into two sub-sets of trials: ‘Congruent’ and ‘Incongruent’. Trials were categorized as Congruent if the f0 and spectral centroid shifted in the same direction. For instance, if the f0 was higher in the second note than the first, so was the spectral centroid. Trials were categorized as Incongruent if the f0 and spectral centroid moved in opposite directions.

To determine the spectral centroid of instruments we first ran instrument notes through STRAIGHT and resynthesized them to be pitch-flattened and harmonic (to match the stimuli used in the experiment), and normalized them to have an rms level of .05. We then estimated the power spectral density using Welch’s method. We used the default parameters of the MATLAB implementation of this method to obtain an estimate of the power at each frequency (1422 samples per window, 50% overlap, defined using Hamming windows with 42.5 dB sidelobe attenuation). We expressed the frequencies in Equivalent Rectangular Bandwidths (ERBs) using the formula of Glasberg and Moore (Glasberg & Moore, 1990), then to estimate the spectral centroid we took a weighted average of these frequencies. The weights were the power (in dB) of each frequency relative to the noise floor, which we estimated by eye to be −65 dB, with values below the noise floor set to 0 so that inaudible frequencies would not contribute to the weighting. We repeated this procedure for 12 different notes (covering a full octave) for each instrument and averaged together the 12 estimates (motivated by the assumption that the spectral envelope was largely the same for all notes, being determined by the instrument body) to obtain an average spectral centroid for each instrument: (from lowest to highest) Ukulele, Pipe organ, Cello, Oboe, and Baritone Saxophone. We note that we did not perform an analogous analysis for Experiment 1, because spectral envelopes in speech are less well summarized by their centroid (because of the salience of multiple formants).

For each of the 20 pre-generated stimulus sets used in the online experiment (described above in Section 2.2), we classified each trial as either Congruent or Incongruent, and then separately analyzed the two sets of trials. Since the Different instrument pairs were randomly selected on each trial (because the experiment was not originally designed to accommodate the congruency analysis), the Congruent and Incongruent trials were not exactly balanced (overall, 51.4% of trials were classified as Congruent, and 48.6% of trials were classified as Incongruent). Since spectral envelopes are often largely determined by the instrument body, in which case they should be independent of the note f0, we categorized trials as Congruent or Incongruent based solely on instrument labels, rather than specifically analyzing each trial.

As observed in the right panel of Figure 3c, performance in the Incongruent condition was significantly worse than that in the Congruent condition (F(1,85)=199.63, *p*<.0001, η_p_^2^=.70). This analysis suggests that the spectral envelope of natural sounds biases pitch discrimination judgments.

### 3.3 Effects of Musicianship

It was a priori unclear whether any of the effects we were testing in Experiments 1 and 2 could be influenced by musical training. Western musical training has been associated with lower pitch discrimination thresholds (Bianchi et al., 2016; Kishon-Rabin et al., 2001; McDermott et al., 2010; McPherson & McDermott, 2018; Micheyl et al., 2006; Spiegel & Watson, 1984). It seemed plausible that musicians (defined here as those with four or more years of musical training) might have more experience comparing notes across different instruments than non-musicians, and thus might be more robust to spectral envelope differences. However, we found no evidence for such differences in robustness. While there was a main effect of musicianship in both experiments (two-way ANOVAs, Experiment 1a: F(1,80)=16.16, *p*=.0001, η_p_^2^=.17 Experiment 2: F(1,84)=8.43 *p*=.0047, η_p_^2^=.09), driven by better discrimination performance in musicians, we did not observe significant interactions between musicianship and same/different vowels or instruments (mixed model ANOVAs, Experiment 1a, Vowels: F(1,80)=1.93 *p*=.18, η_p_^2^=.02, Experiment 2, Instruments: F(1,84)=0.51 *p*=.47, η_p_^2^=.006). There was likewise no interaction between musicianship and harmonicity (mixed model ANOVAs, Experiment 1a, Vowels: F(1,80)=1.88, *p*=.18, η_p_^2^=.02), Experiment 2, Instruments: F(1,84)=0.88, *p*=.35, η_p_^2^=.01). We repeated these analyses using some alternative cutoffs and found that the group differences did not depend sensitively on the cutoff choice. These findings suggest that the effects of spectral envelope variation on pitch perception are not strongly dependent on formal musical training. Given the lack of a musicianship effect in these two experiments, we did not analyze musicianship in subsequent experiments, or in Experiment 1b.

### 3.4 Experiment 3: Discrimination of synthetic tones with extreme spectral envelope differences

#### 3.4.1 Purpose and procedure

Experiment 3 was designed to further explore the biasing effects observed in Experiment 2. We aimed to determine whether the bias would remain in a setting where listeners were forced to completely rely on representations of f0 to complete the task (Figure 4a). To address this question, we tested participants with synthetic tones whose harmonics were completely non-overlapping from note-to-note (as if the result of unnaturally steep rectangular filters that completely attenuate harmonics outside their passband). The filters that produce most natural sounds, as in the instrument tones in Experiment 2, tend to modestly attenuate harmonics rather than render them inaudible. This modest attenuation leaves harmonics in common between notes, and our results here and elsewhere suggest that in many conditions listeners compare these frequencies to detect pitch differences, irrespective of whether notes are harmonic or inharmonic (McPherson et al., 2022; McPherson & McDermott, 2018; McPherson & McDermott, 2020). However, when the two tones contain entirely separate subsets of the harmonic series, discrimination should be impossible when tones are altered to be inharmonic, as there are no common frequency partials to compare. Above-chance performance when tones are harmonic thus requires a representation of the f0. We asked whether listeners are biased by the timbre in such conditions, by testing listeners using notes whose spectral envelopes differed to be either ‘Congruent’ or ‘Incongruent’ with the f0 shift, comparable to Experiment 2.

**Figure 4:**
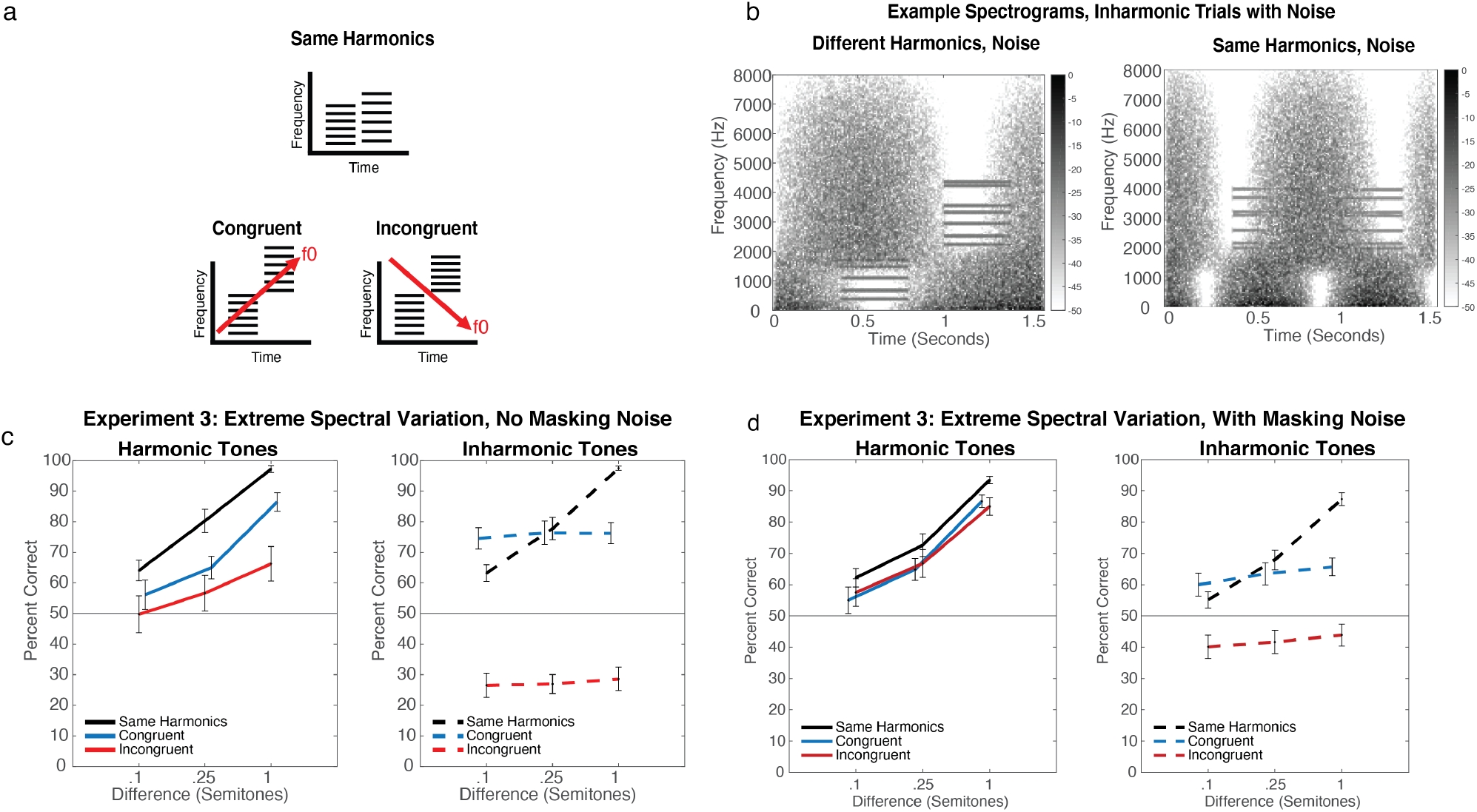
Stimuli and results for Experiment 3, Discrimination of synthetic tones with extreme spectral envelope differences. a. Stimulus configurations. The two tones on a trial could either have the same set of harmonics, or nonoverlapping sets of harmonics, arranged so that the change in spectral centroid was either Congruent or Incongruent with the change in f0. b. Results from Experiment 3, tones presented without noise, grouped by harmonicity. Error bars denote standard error of the mean. c. Example spectrograms of stimuli from an Inharmonic trial with different harmonics (left) or the same harmonics (right), with added masking noise. d. Results from Experiment 4, tones presented with noise, grouped by harmonicity conditions. Error bars denote standard error of the mean.

In half of the trials, tones were embedded in masking noise (Figure 4b). This noise was intended to mask any distortion products that might in principle provide corresponding harmonics between two tones that would otherwise not have them. If participants use distortion products in conditions without noise, the addition of masking noise should impair performance.

As in Experiments 1 and 2, participants heard two tones and were asked whether the second tone was higher or lower than the first tone. Participants completed 30 trials per condition for all Same Harmonic conditions. We had originally intended to average together ‘Congruent’ and ‘Incongruent’ trials in a ‘Different Harmonics’ condition, so participants only completed 15 Congruent trials and 15 Incongruent trials. Congruent and Incongruent conditions were randomly intermixed (randomized independently for each participant).

Psychtoolbox for MATLAB (Kleiner et al., 2007) was used to play sound waveforms. Sounds were presented to participants at 70 dB SPL over Sennheiser HD280 headphones (circumaural) in a soundproof booth (Industrial Acoustics). Participants were paid a fixed hourly rate for their participation. All participants self-reported normal hearing.

#### 3.4.2 Stimuli

The two tones on each trial contained sets of equal-amplitude harmonics, added in sine phase. For ‘Congruent’ and ‘Incongruent’ conditions, one tone contained harmonics 1-5, or 1-6 (low harmonics, chosen equiprobably), and the other note contained harmonics 6-12 or 7-12, respectively (high harmonics, chosen equiprobably) (Figure 4a). In Congruent conditions if the second tone’s f0 (or nominal f0, for inharmonic conditions) was higher than the first, the low harmonics were used in the first note and the high harmonics were used in the second note, and vice versa when the second note’s f0 was lower than the first. Incongruent conditions had the inverse relationship between the f0 and the spectra. For ‘Same Harmonics’ conditions, the same set of high or low harmonics (equiprobably) were used in each tone. The tones were 400 ms in duration, windowed with 20 ms half-Hanning windows, and separated by a 200 ms silent interval.

In half of the trials, tones were presented in quiet, and in the other half, the tones were presented in noise. The noise conditions had originally been intended to look at the possible effect of spectral completion (McDermott & Oxenham, 2008) on the bias induced by the spectral envelope, and so the noise was designed with this goal in mind. We include these conditions here as control conditions to test whether distortion products might underlie the results. The complicated construction of the noise reflects the fact that it was not originally intended to be used exclusively for this purpose. The noise consisted of a superposition of low-pass and high-pass filtered pink noises that were sinusoidally amplitude modulated (100% modulation depth), with phases and modulation frequencies of the modulator waveform hand-chosen to plausibly mask the ‘missing’ harmonics in each tone (Figure 4b). There were 16 possible modulation frequency/phase combinations for the low-pass noise and 16 possible combinations for the high-pass noise (see Supplementary Table 3 for combinations). We used modulated rather than unmodulated noise in an attempt to avoid a direction cue that could plausibly interfere with the discrimination task, potentially obscuring an otherwise beneficial effect of the noise. The low-pass noise was filtered using a sigmoidal transfer function in the frequency domain with an inflection point located two harmonics higher than the highest harmonic of the ‘low’ complex (e.g. the 9^th^ harmonic if the ‘low complex’ contained harmonics 1-7), a slope yielding 40 dB of gain per octave, and a maximum value of 0 dB. The low edge of the high-pass filter was sigmoidal with an inflection point located two harmonics lower than the lowest harmonic of the ‘high’ complex, again with a slope yielding 40 dB of gain per octave and a maximum value of 0 dB.

The same noise filtering parameters were used for the Same Harmonics condition, but different modulation frequencies and phases were used, as we needed to mask the same frequency range for both tones. We hand-selected 5 modulation frequencies and phases to mask the lower frequency regions, and 5 to mask higher frequency regions (see Supplementary Table 4 for exact phase and frequency details). These noise conditions were likewise chosen so that they would fully mask the ‘missing’ harmonics. The purpose of the Same Harmonics conditions was to assess the effects of noise in a situation where it would not cause the spectrum of the tones to be inferred to be more similar than they would be without the noise.

The noise level was set to be high enough in level to mask the missing harmonics were they present, and thus was also sufficient to mask any distortion products generated by the stimulus harmonics (Norman-Haignere & McDermott, 2016; Pressnitzer & Patterson, 2001). The first author conducted an informal adjustment experiment in which she listened to the superposition of the noise and the harmonics that were missing from the main experimental stimuli and iteratively adjusted the noise level to reliably mask the harmonics. This experiment was performed for each of the 16 different masking noises. We settled on a noise level for which the original pink noise signal (prior to filtering and amplitude modulation) was 28 dB higher in overall level than a single harmonic of the tone. Because of the complexity of the noise signal this is the simplest way to report the signal-to-noise ratio to enable replication. The lowpass noise, highpass noise, and tone were added together and the overall level of the combined signal was set to 70 dB SPL. The sampling rate was 16 kHz.

For all conditions, notes could be either harmonic or inharmonic. All inharmonic tones were made inharmonic as described above in *Inharmonic Stimuli*. The f0 (or nominal f0, for inharmonic conditions) of the first tone of each trial was randomly selected from a log uniform distribution spanning 200 to 400 Hz. The f0 of the second tone was either higher or lower (equiprobably) than that of the first tone by the necessary step size: .1, .25, or 1 semitones.

#### 3.4.3 Participants

Experiment 3 was completed in lab. All participants were native English speakers residing in the Greater Boston area. 27 participants completed the experiment but 3 performed below 55% correct across all conditions, and were excluded from further analysis. The data presented here is from the remaining 24 participants (11 self-identified as female, 13 as male, 0 as non-binary; mean age = 32 years, S.D.=14 years). 13 participants had over 4 years of musical training (self-reported, mean = 17, S.D.=13). This sample size allows a 90% chance of seeing a small-to-moderate effect size (Cohen’s f = .2) at a *p*<.05 level of significance.

#### 3.4.4 Results and discussion

Although performance was above chance in all harmonic tone conditions, we found that the impairment observed in Experiment 2 persisted for synthetic tones with non-overlapping harmonics (Figure 4c, left). Performance was better when listeners heard the same harmonics from note-to-note than in both Congruent and Incongruent conditions (Same Harmonics vs. Congruent, Harmonic tones, F(1,23)=7.26, *p*=0.013, η_p_^2^=.24; Same Harmonics vs. Incongruent, Harmonic tones, F(1,23)=21.51, *p*=0.0001, η_p_^2^=.48). While there was not the significant main effect of congruency that would signify a spectral bias (F(1,23)=2.06, *p*=0.16, η_p_^2^=.08), this appears to be due to near-chance performance for the smaller step sizes. At the 1 semitone step size, performance in the Incongruent condition was significantly worse than that for the Congruent condition (t(23)=2.15, *p*=.005). The spectral bias observed in Experiment 2 thus appears present in these conditions as well.

As in previous experiments with synthetic tones (Faulkner, 1985; McPherson & McDermott, 2018; McPherson & McDermott, 2020; Micheyl et al., 2010; Moore & Glasberg, 1990), and consistent with Experiment 1, performance was similar for Harmonic and Inharmonic tones when the two tones had the same harmonics (Figure 4c, black solid and dashed lines, F(1,23)=0.65, *p*=0.43, η_p_^2^=.03). However, as expected, listeners could not perform the task with Inharmonic tones whose harmonics did not overlap, with performance entirely driven by the direction of spectral centroid change: there was no main effect of step size (F(1,23)=1.00, *p*=.38, η_p_^2^=.04). This suggests that – as intended based on the design of the tones – listeners rely on the f0 when notes are harmonic and spectral envelope differences are sufficiently extreme. But even in this case, spectral invariance is incomplete.

The conditions with added noise help to rule out the possibility that discrimination in the different harmonics conditions was mediated by distortion products rather than the inferred f0, because performance in the Congruent conditions was similar with and without noise (F(1,23)=0.017, *p*=0.90, η_p_^2^=.0007), and in the Incongruent conditions was actually better with noise (F(1,23)=13.55, *p*=0.0012, η_p_^2^=.37, Figure 4d). Given that the noise was sufficient to mask any distortion products (Norman-Haignere & McDermott, 2016; Pressnitzer & Patterson, 2001), it should have impaired performance (and certainly not improved it) if distortion products were supporting discrimination.

The masking noise we added also created a scenario in which the two sounds to be compared could have similar spectral envelopes, with different parts of their spectra obscured by masking noise. In such settings the auditory system is known to “fill in” masked portions of the spectrum (McDermott & Oxenham, 2008). The fact that masking noise caused pitch judgments to become more robust to spectral envelope variation between sounds is consistent with the idea that the noise induces spectral completion on the part of the listeners, and that the spectral bias is determined by the inferred spectral envelope of the tones.

### 3.5 Experiment 4: Effect of time delay on spectral invariance

#### 3.5.1 Purpose and procedure

Experiment 3 showed that pitch discrimination judgments are biased by the spectral envelope of notes even in conditions where listeners must rely on the f0 to perform the task because the harmonics are completely non-overlapping. Two distinct hypotheses could explain this bias. One possibility is that the representation of a sound’s f0 is biased by the sound’s spectral envelope. Another possibility is that the f0 representation is itself unbiased, but that listeners are unable to base decisions exclusively on this representation, with spectral envelope differences between notes biasing their judgments.

In Experiment 4 we attempted to disambiguate these hypotheses using time delays. In previous work we found that listeners become more reliant on representations of the f0 when making judgments about pitch across a delay (McPherson & McDermott, 2020), as though the f0 representation is better remembered than representations of the individual frequencies. Although not tested in previous work, it seemed plausible that the f0 might also be better remembered than the spectral envelope. If the f0 representation is itself unbiased, with spectral biases coming from suboptimal use of cues at a decision stage, it seemed possible that the bias would be reduced over a delay.

As in Experiment 3, tones either contained the same set of harmonics or non-overlapping harmonics. On the non-overlapping trials, the spectral envelope could be either Congruent or Incongruent with the f0 change (Figure 5a). Listeners heard two notes during each trial, either played back-to-back or separated by a 3 second delay (Figure 5b). We only tested participants using Harmonic notes. The task was always to say whether the second tone was higher or lower than the first, and participants received feedback (Correct/Incorrect) after each trial. Participants completed 24 trials per condition.

**Figure 5:**
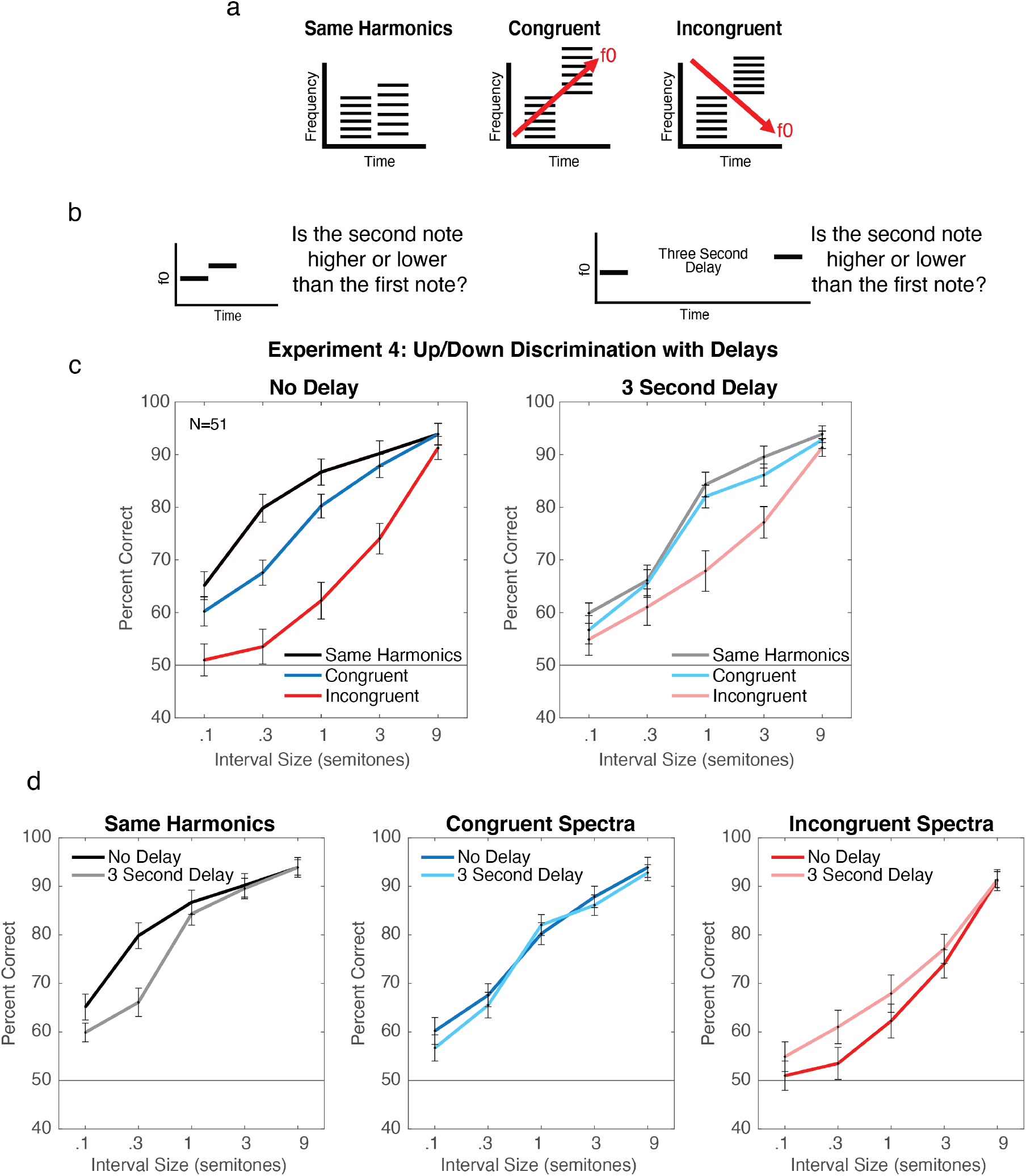
Stimuli and Results for Experiment 4, Effect of delays on discrimination performance with synthetic tones. a. Stimulus configurations. The two tones on a trial could either have the same set of harmonics, or nonoverlapping sets of harmonics, arranged so that the change in spectral centroid was either Congruent or Incongruent with the change in f0. Tones could either be presented back-to-back (no delay; top row), or separated by a three second delay (bottom row) b. Task for all experiments. Participants heard two sounds and judged which was higher in pitch. c. Results from Experiment 4, grouped by delay conditions. Here and in c, error bars denote standard error of the mean. d. Results from Experiment 4, grouped by spectra condition.

#### 3.5.2 Stimuli

Tones were identical to those used in Experiment 3, except they were either played back-to-back, or separated by a 3 second delay. The f0 of the first tone of each trial was randomly selected from a log uniform distribution spanning 200 to 400Hz.

#### 3.5.3 Participants

194 participants passed the headphone check and completed Experiment 4 online. 87 had overall performance (averaged across all conditions) below 55% correct and were removed from further analysis. The large number of participants excluded based on this criterion partly reflects the fact that data was collected during a time when Amazon’s Mechanical Turk platform was experiencing a higher-than-normal level of fraud, and we observed that many participants were performing at chance levels. Of the 107 remaining participants, 39 identified as female, 67 as male, 1 as non-binary. The average age of these participants was 37.2 years (S.D.=10.0). 35 participants had four or more years of musical training, with an average of 10.6 years (S.D.=10.1).

To determine sample size, we performed a power analysis by bootstrapping pilot data (from an earlier version of Experiment 4 in which listeners heard all but the 9-semitone step size condition). For each possible sample size, we computed bootstrap distributions of the interaction F statistic (Congruent/Incongruent vs. No Delay/Delay). We found that a sample size of 11 yielded a 95% chance of seeing the interaction present in our pilot data at *p*<.01 significance level. However, based on our pilot data we also wanted to have sufficient power to detect an effect of delay specifically on the Incongruent condition (where we found that performance improved with delay). A power analysis using G*Power (Faul et al., 2007) indicated that we needed at least 68 participants to be 95% sure of seeing an effect half of that observed in the pilot study (η_p_^2^=.13) at a *p*=.01 significance level. In practice, we ran the experiment in small batches and ended up with more than this number of participants.

#### 3.5.4 Results and discussion

As shown in Figure 5c, the effects of the spectral envelope decreased with delay, producing a significant interaction between Congruent/Incongruent conditions and Delay (F(1,106)=21.89, *p*<.0001, η_p_^2^=.17). We replicated the pronounced effect of spectral envelope variation observed in Experiment 3 when there was no delay (performance was worse for Congruent than for Same Harmonics (F(1,106)=8.13, *p*=.005, η_p_^2^=.07, and worse for Incongruent than Congruent (F(1,106)=48.95, *p*<.0001, η_p_^2^=.32). However, these effects were reduced when there was a delay between tones, due to the delay having different effects in different stimulus conditions. The delay reduced performance for the Same Harmonics condition (Figure 5d, significant main effect of delay; F(1,106)=17.48, *p*<.0001, η_p_^2^=.14). But there was no significant effect of delay in the Congruent Spectra condition (F(1,106)=2.48, *p*=.12, η_p_^2^=.02). And the delay in fact *improved* performance in the Incongruent condition (significant effect of delay, but in the opposite direction; F(1,106)=15.62, *p*=.0001, η_p_^2^=.13).

It is not obvious how to account for these results without positing that the representations of pitch and spectral envelope are separate. One might suppose instead that the effect of the delay could reflect strategies such as subvocal or active rehearsal of the stimulus that could serve to reduce bias, but several previous experiments have suggested that active rehearsal strategies do not aid pitch discrimination in such settings (Kaernbach & Schlemmer, 2008; Massaro, 1970; McPherson & McDermott, 2020). And if the representation of pitch were itself biased, the bias should persist across delays, with the delay making performance worse for both Congruent and Incongruent conditions. In this respect the most diagnostic result is the counterintuitive improvement with delay in the Incongruent condition. The change in bias with delay suggests that the representations of pitch and spectral envelope are likely independent, and decay at different rates, with the effect of the spectral envelope on pitch judgments decreasing over time.

### 3.6 Experiment 5: Spectral invariance of musical interval perception

#### 3.6.1 Purpose and procedure

Experiment 5 was intended to test whether the spectral bias we observed across several pitch discrimination tasks (Experiments 1-4) would generalize to judgments about the magnitude of pitch differences (intervals). This question was previously addressed by Russo and Thompson, who asked listeners to rate interval sizes on a scale of 1 to 5, and found that timbre changes could augment or diminish the rated interval size (Russo & Thompson, 2005). The purpose of our experiment was to test whether this effect would occur for a musical judgment based on pitch intervals. In our experiment, listeners heard two notes and were asked whether the notes matched the starting interval in ‘Happy Birthday’, specifically, the two-semitone difference between the last syllable of ‘Happy’ and the first syllable of ‘Birthday’ (Figure 6a). Listeners either heard notes separated by the correct interval of 2 semitones, or by an interval that was mistuned by up to a semitone in either direction (2 semitones +/- .5 or 1 semitones). Participants completed 6 trials per condition.

**Figure 6:**
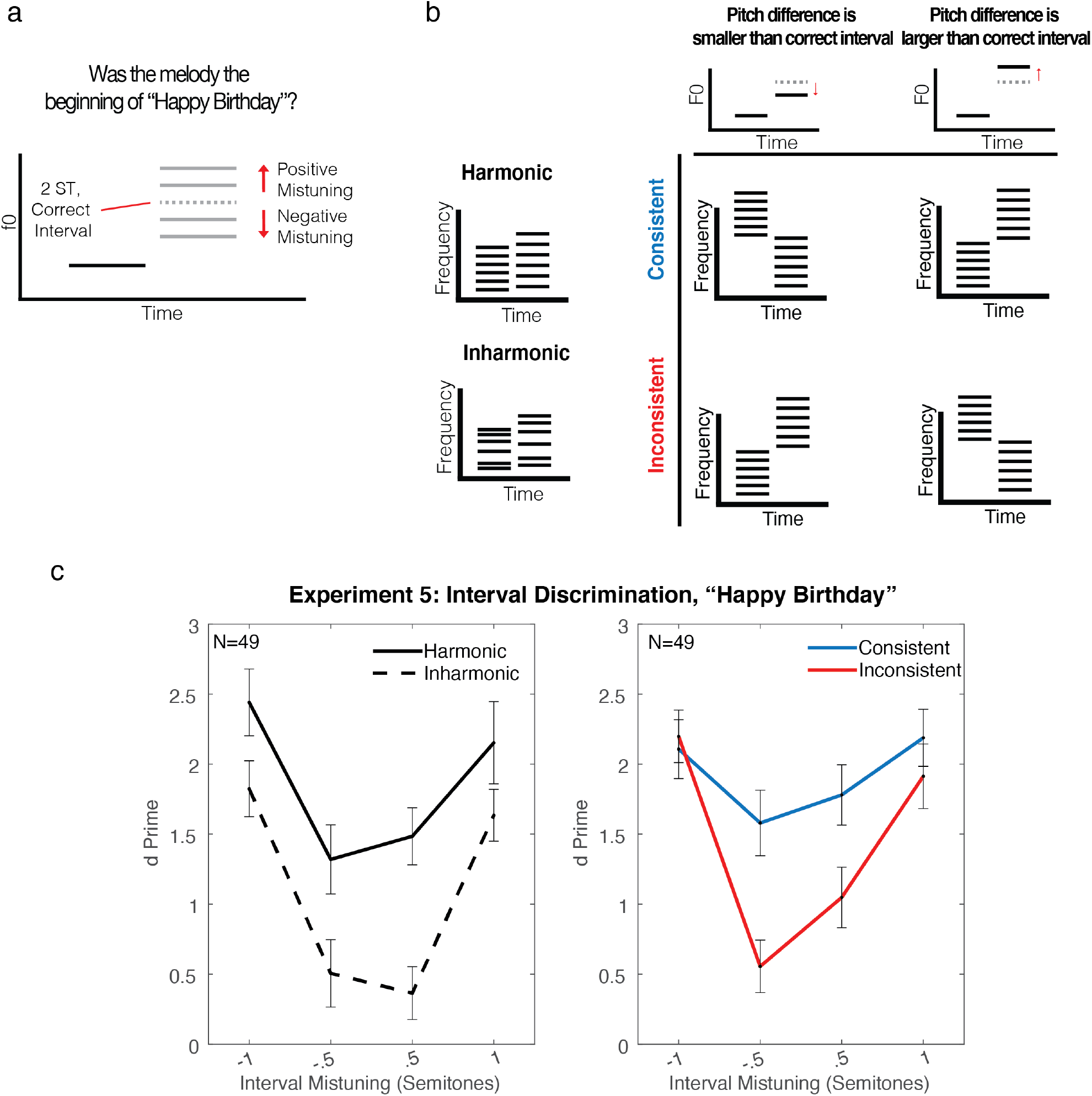
Stimuli and Results for Experiment 5, Discrimination of intervals composed of synthetic tones with extreme spectral envelope differences. a. Schematics of a trial. b. Schematics of stimulus conditions. Two right-most columns show the relationship between the spectra and f0 in ‘Consistent’ and ‘Inconsistent’ conditions. c. Results from Experiment 5. Error bars denote standard error of the mean.

We tested four conditions: Harmonic, Inharmonic, Consistent, and Inconsistent (Figure 6b). The Harmonic and Inharmonic conditions presented pairs of tones identical to those in Experiment 3 apart from the different pitch intervals used. The purpose of these conditions was to establish baseline performance on the task, and to help characterize the representations used for the task (namely, whether performance was dependent on representations of the f0). The latter seemed advisable given that this was not a task we had previously used.

The ‘Consistent’ and ‘Inconsistent’ conditions were analogous to the ‘Congruent’ and ‘Incongruent’ conditions in Experiments 3 and 4. However, the second note in each trial was always higher than the first note, so instead of the change in spectral centroid being aligned or mis-aligned with the f0 change, it was aligned or mis-aligned with the positive or negative mistuning from the correct two-semitone interval. For example, in the ‘Consistent’ condition, if the pitch mistuning was positive (the presented interval was larger than two semitones), then the first note contained the lower set of harmonics and the second note contained the higher set of harmonics, and vice versa for a negative mistuning. Our prediction was that the spectral shift might increase or decrease a listener’s estimate of the interval size, making the correct answer more apparent in the ‘Consistent’ case, and less apparent in the ‘Inconsistent’ case.

#### 3.6.2 Stimuli

On each trial participants were presented with two synthetic complex tones in succession (with no delay between tones). Tones were identical to those used in Experiment 3. The f0 (or nominal f0, for Inharmonic conditions) of the first tone of each trial was randomly selected from a log-uniform distribution spanning 200 to 400 Hz. The f0 of the second tone was either 2 semitones above the first (to match the first interval in ‘Happy Birthday’ or mistuned from 2 semitones by +/- .5 or 1 semitone. On Consistent and Inconsistent trials the spectral centroid of the notes moved in the same or opposite direction as the mistuning, as described above. There were an equal number of ‘Happy Birthday’ and ‘Not Happy Birthday’ trials throughout the experiment.

#### 3.6.3 Participants

252 participants passed the headphone check and completed Experiment 3 online. 203 had performance (averaged across all conditions) below a d-prime of .67 and were removed from further analysis. We chose this exclusion criteria based on a small pilot study completed in the lab before the COVID-19 pandemic, with 5 participants completing the task with Harmonic and Inharmonic tones. Across harmonicity conditions, the average d-prime across the two smallest mistunings (+/-.5 semitones) was .67, so we used this as a conservative estimate of good performance among online participants. As in Experiment 4, the large number of participants excluded based on this criterion partly reflects the fact that data was collected during a period when Amazon’s Mechanical Turk platform was experiencing a higher-than-normal level of fraud, and we observed that many participants were performing at chance levels. Of the remaining 49 participants, 23 identified as female, 24 as male, 2 as non-binary. The average age of these participants was 38.5 years (S.D.=11.5). 23 participants had four or more years of musical training, with an average of 14.6 years (S.D.=11.7).

The target sample size was determined based on a separate pilot experiment, run online, where participants only heard Harmonic and Inharmonic conditions. In this experiment participants heard 10 trials per condition, rather than 6. To estimate an effect size with fewer trials per participant we bootstrapped from pilot data, sub-sampling 6 trials from each participant per condition and calculated the Harmonic vs. Inharmonic effect size 10,000 times. The average effect size was η_p_^2^=.23. We predicted that effects of the spectrum might be smaller than effects of harmonicity. A power analysis indicated that we would need 38 participants to observe an effect 1/2 the size of the effect of harmonicity at a p value of .01, 95% of the time (Faul et al., 2007).

#### 3.6.4 Results and discussion

Participants were well above chance in the Harmonic condition, with performance improving with the magnitude of the mistuning, as expected. Performance remained above chance in the Inharmonic condition, but was substantially worse than in the Harmonic condition (Figure 6c, F(1,48)=17.86, *p*=.0001, η_p_^2^=.27). These results complement previous findings that tasks related to musical interval perception are impaired with inharmonic tones, thus likely relying on f0-based pitch (McPherson & McDermott, 2018). The above-chance performance with inharmonic tones suggests that listeners can nonetheless extract some information about interval size from shifts in the partials.

Parallel to the spectral biases found for pitch discrimination in Experiments 1-4, performance was better in the Consistent condition compared to the Inconsistent condition (F(1,48)=6.94, *p*=.011, η_p_^2^=.13). This suggests that the limited invariance of pitch judgments is not specific to up-down pitch discrimination. This result is also consistent with the idea that the bias in pitch judgments originates in representations of relative pitch, rather than exclusively at a decision stage. Unlike a pitch discrimination task, here the decision involves judging whether the heard interval is the same or different as a remembered interval. It is not obvious how the sense that the timbre went up or down would interfere with the decision to respond “same” or “different”. Rather, the timbre change seems to be incorporated into the pitch interval estimate. Relative pitch representations may thus be biased by changes in the spectral envelope, even if the f0 representation they are based on is not. These biases in relative pitch could also explain the effects of spectral envelope seen on pitch discrimination (Experiments 1-4).

## 4 GENERAL DISCUSSION

We tested the extent to which listeners can make relative pitch judgments about sounds differing in their spectral envelopes, both for natural sounds such as instruments and voices, and synthetic tones with extreme levels of spectral envelope variation. Our first question was whether any spectral invariance in pitch perception is mediated by representations of f0, in which case discrimination of sounds differing in spectral envelope should be worse when the sounds are inharmonic. We observed inharmonic deficits when f0 differences between otherwise natural sounds were large, but these deficits were similar for conditions where the spectral envelope (vowel or instrument) was the same vs. different. The only cases in which spectral envelope variation produced impairments specific to inharmonic stimuli involved synthetic tones with extreme spectral envelope differences between notes. Our second question was whether the limits of invariance reflect biases in representations underlying pitch vs. processes that operate on those representations to mediate pitch comparisons. We found that the spectral bias of pitch judgments decreased across a delay, suggesting that the bias is not exerted on the representation of a sound’s f0. Taken together, our results show that human pitch perception exhibits some degree of spectral invariance, enabling pitch comparisons between natural sounds with different spectral envelopes, but that this invariance does not normally require representations of f0, or make use of the fact that f0 representations are apparently unbiased.

### 4.1 Real-world pitch discrimination is somewhat robust to timbre, independent of f0-based pitch

We built on previous studies examining pitch discrimination between tones differing in spectral envelope (Allen & Oxenham, 2014; Chuang & Wang, 1978; Hellström et al., 1994; Kuhl & Miller, 1982; Micheyl & Oxenham, 2004; Miller, 1978; Moore et al., 1992; Moore & Glasberg, 1990; Repp & Lin, 1990; Russo & Thompson, 2005; Siedenburg et al., 2022; Singh & Hirsh, 1992; Stoll, 1984; Warrier & Zatorre, 2002), by using comparisons between harmonic and inharmonic stimuli to test whether invariances in pitch perception are dependent on f0-based pitch. In both Experiment 1 (speech) and Experiment 2 (musical instruments), participants showed above-chance up-down discrimination irrespective of whether the instruments or speakers being compared were the same or different. This result confirms that pitch perception is somewhat robust to everyday spectral envelope differences between sounds, but is nonetheless worse when different instruments or vowels were compared. However, the impairments resulting from between-sound spectral envelope variation were no greater for inharmonic sounds. In particular, for small f0 differences (3 semitones and lower) discrimination performance was similar for harmonic and inharmonic conditions. The results are consistent with other evidence that listeners do not normally use f0 representations when discriminating sounds presented back-to-back with modest f0 differences (McPherson et al., 2022; McPherson & McDermott, 2018; McPherson & McDermott, 2020), but suggest that this is the case even for spectral envelope differences between sounds typically encountered in natural conditions. For larger f0 differences discrimination was impaired for inharmonic tones, but these effects held regardless of whether the instruments or vowels were the same or different. This result suggests that human robustness to everyday spectral envelope variation does not rely on representations of the f0.

### 4.2 Invariance to extreme spectral envelope differences is mediated by f0-based pitch

In Experiment 3, we introduced extreme (and unnatural) spectral envelope changes between notes by presenting nonoverlapping subsets of frequency partials. In these conditions, listeners were by design only able to make accurate discrimination judgments when the stimuli were harmonic, indicating a reliance on representations of f0. However, performance with harmonic tones was nonetheless worse when the two notes did not share harmonics. Both findings are consistent with previous studies examining pitch discrimination between sounds that vary in their spectra (Allen & Oxenham, 2014; Micheyl & Oxenham, 2004; Moore et al., 1992; Moore & Glasberg, 1990; Singh & Hirsh, 1992; Warrier & Zatorre, 2002). It is thus clear that listeners can use representations of the f0 when there is extreme spectral envelope variation between sounds, but this paper provides evidence that such experimental manipulations are not representative of what happens when we discriminate natural sounds. Evidently the spectral envelope differences between natural sounds are sufficiently modest that any invariance does not require f0-based pitch.

### 4.3 Necessity of f0-based pitch for registering large f0 changes

In Experiments 1 and 2 we found that when f0 changes were sufficiently large, performance with inharmonic stimuli was impaired relative to harmonic stimuli. This impairment plausibly reflects ambiguities in the correspondence between frequency partials due to the filter that is present for most natural sounds. When pitch changes are small the correspondence between frequency partials of two sounds being compared is likely to be unambiguous. This is because the f0 difference is small relative to the harmonic spacing, such that a resolved frequency partial in the first sound has a close match at the corresponding harmonic in the second sound. However, when pitch changes become sufficiently large, the correspondence between partials may become ambiguous when a filter attenuates the upper and lower ends of the source spectrum (Figure 2f). This correspondence could be aided by some form of integration across frequency (to find the best match collectively for all of the partials associated with a source), but f0-based pitch evidently also helps listeners to register such large changes in f0, producing improved discrimination for harmonic tones. In a previous study we observed analogous effects with synthetic tones passed through a fixed bandpass filter (replotted in Supplementary Figure 1 for comparison). The contribution of the present experiments is to show that such effects occur in natural sounds and thus have relevance for real-world listening. The inharmonic deficit at large intervals was present for both speech and music sounds, providing some evidence for a domain-general phenomenon.

### 4.4 Origins of timbral biases in pitch judgments

The results of Experiments 1-3 left it unclear whether observed pitch discrimination biases from spectral envelope variation are the result of bias in the pitch estimation process or bias at some subsequent stage, such as the task decision (up vs. down). We leveraged the effects of time delay on pitch discrimination to explore these possibilities. Previous experiments have suggested that listeners become more reliant on representations of f0 when comparing sounds across a time delay (McPherson & McDermott, 2020), potentially because representations of the constituent frequency partials of a sound deteriorate more rapidly over time than those of the f0. This result raised the possibility that representations of the spectral envelope might also deteriorate more rapidly than those of the f0, in which case time delays might help to distinguish the origin of the spectral envelope bias. We reasoned that if the bias occurs at a stage where the pitch of different sounds is being compared, and if spectral envelope information decays more over time than does information about the f0, then the observed bias might change with the delay. Experiment 4 substantiated this prediction – the bias in discrimination judgments decreased across a delay, and in particular, performance improved with an inter-note delay for trials where the spectral envelope and f0 were incongruent. One speculative possibility is that f0-based pitch is remembered better than timbre due to the need to store pitch over relatively long temporal extents for use in music cognition. Regardless of the root cause, these results suggest that the bias in pitch judgments is likely not the result of bias in the representation of the f0 of individual sounds, but rather arises at a subsequent comparison stage. This conclusion is consistent with findings that pitch matching is not much affected by note timbre (Russo & Thompson, 2005).

Experiment 5 provided evidence that spectral biases occur at the stage of relative pitch representations, i.e. representations of pitch changes. The task in that experiment required listeners to compare the stimulus to their memory of the first interval in ‘Happy Birthday’, and make a same-different judgment. The results suggest that timbre differences bias the representation of pitch intervals, replicating findings of Russo and Thompson (2005) but with a musical judgment. In contrast to the biases seen in pitch discrimination (Experiments 1-4), it is not obvious how to explain the bias in interval judgments as occurring purely at a decision stage. With up-down pitch discrimination judgments, the bias in decisions could result from a competing direction signal from the spectral envelope being difficult for listeners to ignore when they choose “up” or “down”. In principle, such decision-stage interference could also occur with the interval size ratings measured by Russo and Thompson. But in the interval task we used in Experiment 5, the judgment was a same/different judgment in which the stimulus was compared to a memory representation. Moreover, the pitch interval stimulus always had the same direction, and differed only in its magnitude. It is thus less obvious how the direction of the timbral change from note to note could aid or impair the decision process. One possibility is that changes in spectral envelope are integrated with changes in pitch at the stage of relative pitch representations (Russo & Thompson, 2005), which in our experiment could result in augmented or diminished representations of the pitch interval between the two notes. Biases in relative pitch representations could also account for the biases seen in pitch discrimination, on the assumption that the decision variable is a representation of relative pitch (the pitch difference between two sounds). The influence of coarse spectral envelope changes on relative pitch is consistent with findings that humans extract melody-like representations from such changes (Cousineau et al., 2014; Graves et al., 2014; Graves et al., 2019; McDermott et al., 2008; Siedenburg, 2018).

### 4.5 Limitations

Our natural sound stimuli were constrained by available corpora. We attempted to test the upper end of naturally occurring spectral envelope variation by choosing pairs of vowels (Experiment 1) and instruments (Experiment 2) with maximally different excitation patterns. We think it likely that the extent of variation in our stimuli is representative of that encountered in everyday listening, but there could be natural contexts where the spectral envelope variation between sounds exceeds the levels that we tested. It is therefore possible that spectra occasionally vary enough from sound-to-sound as to necessitate f0-based pitch discrimination in real-world conditions. However, this does not appear to be the norm for speech and music sounds, at least for those we had access to here.

Our experiments with natural sounds relied on resynthesis (to render harmonic and inharmonic versions of sounds that were otherwise matched), and this resynthesis might in principle limit the naturalness of the stimuli. The resynthesis is clearly not perfect, but our subjective sense is that the artifacts it induces are modest. Moreover, the extent of the artifacts appears similar for harmonic and inharmonic sounds (the same estimated spectral envelope was used in the resynthesis of harmonic and inharmonic sounds, in part to help minimize any such synthesis differences). Consistent with this observation, speech intelligibility of resynthesized harmonic and inharmonic speech utterances is indistinguishable in quiet (McPherson et al., 2022; Popham et al., 2018). While the synthesis method used in this study was originally designed for speech, the same principles of estimating source vs. filter apply to musical instruments, and our subjective sense is that the timbre of harmonic instruments remained identifiable after resynthesis. Example vowel and instrument stimuli are available at https://mcdermottlab.mit.edu/SpectralVariation.html for readers to judge for themselves.

### 4.6 The function of f0-based pitch

The current results are consistent with a growing body of literature suggesting that in many contexts listeners may use representations of constituent frequency partials, rather than the f0, for “pitch” discrimination (Chambers et al., 2017; Demany & Ramos, 2005; Faulkner, 1985; McPherson et al., 2022; McPherson & McDermott, 2018; McPherson & McDermott, 2020). The results here provide additional evidence that this phenomenon occurs for natural sounds, in particular speech and musical instruments. The key evidence comes from results with inharmonic stimuli – even though the f0 of individual inharmonic sounds is ambiguous, the change in frequency partials from one sound to another is not (at least for modest f0 differences), and listeners evidently hear this signal as a pitch change and rely on it to make judgments. The results suggest that in many real-world contexts, changes in frequency partials accurately convey f0 changes, and are used to make judgments about the f0. We argue that representations of the partials and of the f0 should both be considered part of pitch perception, as both can be used by listeners to make judgments about f0 changes depending on the conditions (McPherson et al., 2022; McPherson & McDermott, 2018; McPherson & McDermott, 2020).

The results here complement evidence that representations of f0 are used to meet at least two other behavioral challenges: retaining information about sounds over time, and hearing in noise. In other studies we found harmonic advantages when the sounds being discriminated were either separated by a time delay (McPherson & McDermott, 2020), or superimposed on noise (McPherson et al., 2022). This study adds a third situation in which harmonic advantages are evident – when there are large f0 differences between sounds passed through a bandpass filter, as is typical for speech and instruments (and many animal vocalizations). By contrast, we found no evidence that f0-based pitch enables invariance to differences in spectral envelope that occur between natural sounds. The reliance on f0 representations in settings with extreme spectral envelope variation (e.g. synthetic tones with distinct sets of harmonics) thus appears to be a byproduct of other functions of f0-based pitch, including compression for memory, noise robustness, and robust estimation of large f0 jumps. These findings clarify our understanding of the functional role of f0 representations.

### 4.7 Future directions

Although we have documented the extent of spectral invariance in pitch perception and clarified the representations that mediate it, we lack a theoretical understanding of the limits to invariance. It is not obvious why human judgments are not more invariant to differences in the spectral envelope of sounds being compared. In particular, listeners do not always adopt strategies that are optimal, at least in the context of certain experimental tasks. For instance, in all experiments we found that discrimination was impaired by spectral envelope differences between sounds, as though listeners are unable to base judgments entirely on the f0 even when that would maximize task performance. It might be that for natural sounds, changes in f0 are strongly correlated with changes in the spectral envelope (Siedenburg et al., 2021), such that the biases we observed are a signature of an optimal strategy for natural sound discrimination. Recent evidence from infants suggests they may be more robust to spectral envelope differences than adult listeners (Lau et al., 2021), raising the possibility that the biases are learned from exposure to natural sounds. Models of pitch discrimination could help to clarify these properties of human pitch perception. Similar questions could be posed about the dependence (or lack thereof) of timbre on pitch (Allen & Oxenham, 2014; Marozeau et al., 2003).

The apparent spectral invariance of f0-based pitch representations, in contrast to the limited invariance of pitch judgments they are assumed to subserve, raises the question of where in the brain these effects arise (Allen et al., 2017; Allen et al., 2022; He & Trainor, 2009; Norman-Haignere et al., 2013; Patterson et al., 2002; Penagos et al., 2004; Tang et al., 2017). Proposed neural correlates of f0 in non-human animals have been reported to be highly invariant to the spectral envelope in some cases (Bendor & Wang, 2005), with some analogous evidence in humans (Allen et al., 2022), and models optimized to estimate f0 in natural sounds produce relatively invariant representations as well (Saddler et al., 2021). However, auditory cortical responses have also been proposed to underlie pitch judgments (Bizley et al., 2013), and might thus be expected to exhibit spectral biases.

## Acknowledgements

The authors thank S. Norman-Haignere and L. Demany for helpful discussions and for comments on an earlier version of the manuscript, and M. Saddler for comments on an earlier version of the manuscript. This work was supported by National Institutes of Health (NIH) grants F31DCO18433 and R01DC014739. The funding agency was not otherwise involved in the research, and any opinions, findings, and conclusions or recommendations expressed in this material are those of the authors and do not necessarily reflect the views of the NIH.

---

**Supplementary Figure 1:**
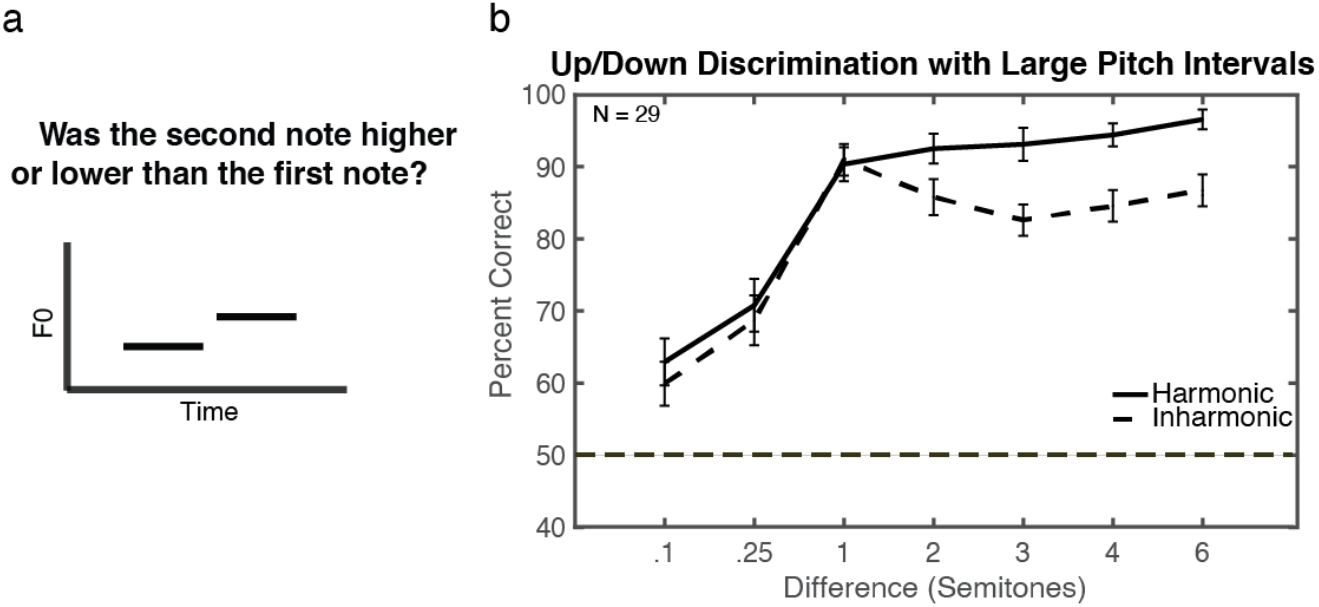
a. Schematics of task and instructions. b. Results from previous study showing large interval effect (McPherson, M. J., & McDermott, J. H. (2018). Diversity in pitch perception revealed by task dependence. *Nature Human Behavior*, *2*, 52-6.). Tones in that study were made inharmonic in the same way that tones were made inharmonic in the current study. The tones in this experiment were filtered with a fixed bandpass filter (Gaussian transfer function on a log-frequency scale, centered at 2,500Hz with a standard deviation of half an octave), applied to ensure that participants could not perform the tasks using changes in the spectral envelope. The tones also had low pass filtered pink noise added to mask distortion products. As in Experiments 1 and 2, with vowels and instrument notes, Harmonic and Inharmonic performance was matched at smaller pitch changes, but performance was impaired for inharmonic tones for larger pitch differences. Error bars show standard error of the mean.

**Supplementary Table 1:**
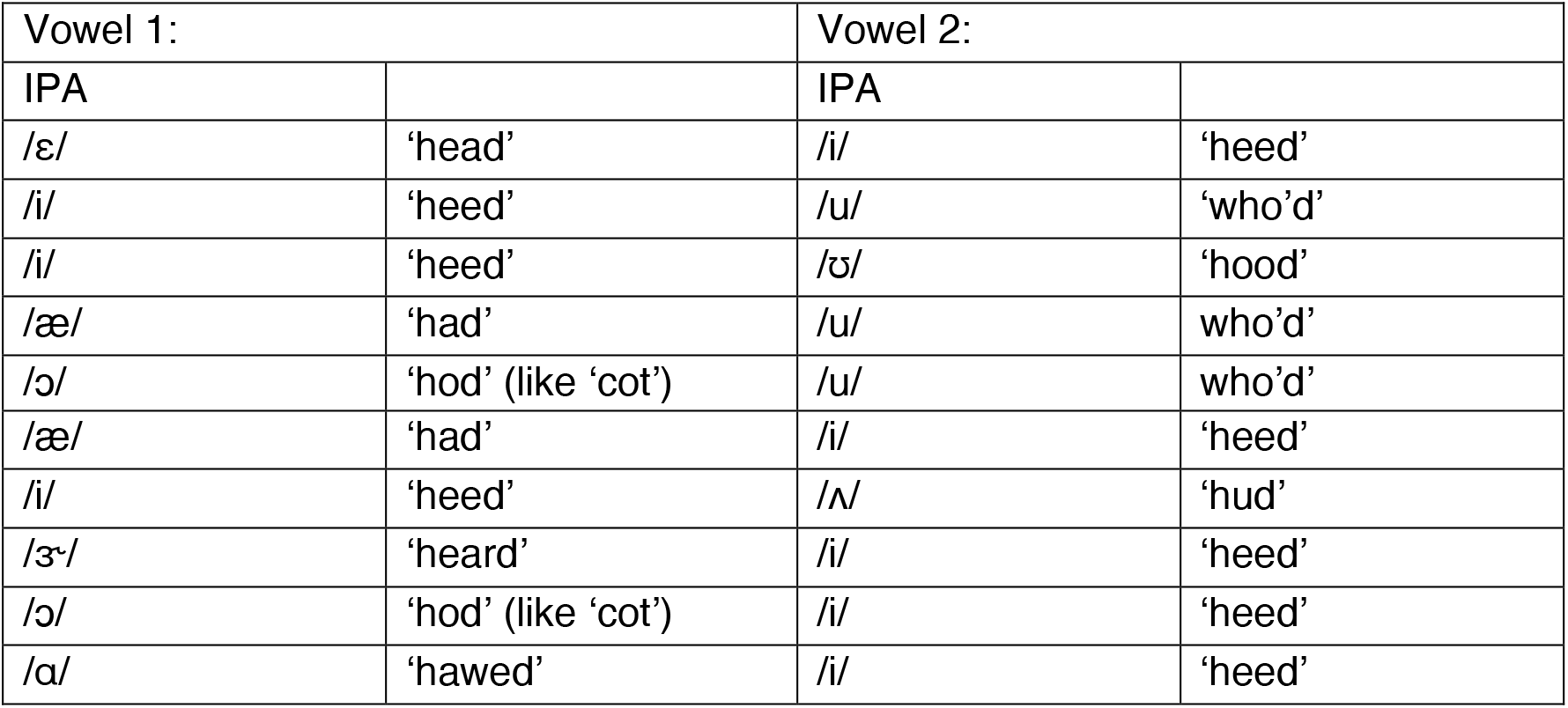
Vowel Pairs (Vowel 1 vs. Vowel 2 order was randomized in the experiment)

**Supplementary Table 2:** Instrument Set

|  |
| --- |
| 1) Alto saxophone |
| 2) Baritone saxophone |
| 3) Bassoon |
| 4) Cello |
| 5) Clarinet |
| 6) Classical guitar |
| 7) Clarinet |
| 8) Cornet |
| 9) Electric guitar |
| 10) English horn |
| 11) Flute |
| 12) French Horn |
| 13) Hammond organ |
| 14) Harpsichord |
| 15) Mandolin |
| 16) Pan flute |
| 17) Piano |
| 18) Pipe organ |
| 19) Soprano saxophone |
| 20) Tenor saxophone |
| 21) Trombone |
| 22) Trumpet |
| 23) Ukulele |
| 24) Viola |
| 25) Violin |

**Supplementary Table 3:** Modulation frequencies and phases for noise amplitude modulation (100% modulation depth)

| Low Pass Noise |  | High Pass Noise |  |
| --- | --- | --- | --- |
| Frequency (Hz) | Phase (radians) | Frequency (Hz) | Phase (radians) |
| 2.5 | $\pi$ | 2.5 | 1 |
| 2 | $2.14 (\pi-1)$ | 2 | 0 |
| $1.\overline{66}$ | $2.14 (\pi-1)$ | $1.\overline{66}$ | -.5 |
| $1.\overline{66}$ | $2.64 (\pi-1.5)$ | $1.\overline{66}$ | -1 |
| 1.429 | 1 | 1.429 | -1 |
| 1.429 | .5 | 1.429 | -1.5 |
| 1.25 | .5 | 1.25 | -2 |
| 1.25 | 0 | 1.25 | -2.5 |
| $1.\overline{11}$ | 0 | $1.\overline{11}$ | -2.5 |
| $1.\overline{11}$ | -.5 | $1.\overline{11}$ | $\pi$ |
| 1 | -.5 | 1 | $\pi$ |
| 1 | -1 | 1 | 2.5 |
| $0.\overline{90}$ | -1 | $0.\overline{90}$ | 2.5 |
| $0.\overline{90}$ | -1.5 | $0.\overline{90}$ | 2 |
| $0.8\overline{3}$ | -2.2 | $0.8\overline{3}$ | 1.5 |
| 0.769 | $\pi$ | 0.769 | 1 |

**Supplementary Table 4:**
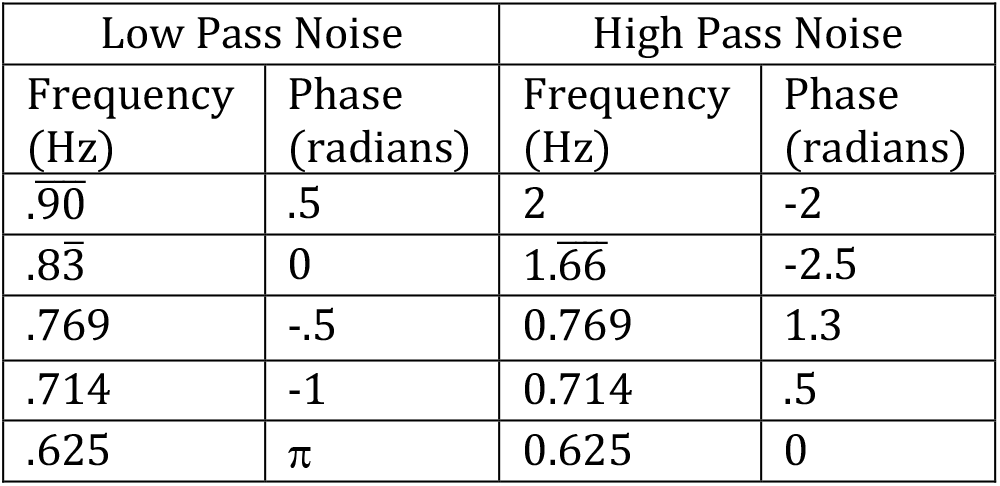
Modulation frequencies and phases for noise amplitude modulation for control conditions (100% modulation depth)

